# SARS-CoV-2 induces double-stranded RNA-mediated innate immune responses in respiratory epithelial derived cells and cardiomyocytes

**DOI:** 10.1101/2020.09.24.312553

**Authors:** Yize Li, David M Renner, Courtney E Comar, Jillian N Whelan, Hanako M Reyes, Fabian Leonardo Cardenas-Diaz, Rachel Truitt, Li Hui Tan, Beihua Dong, Konstantinos Dionysios Alysandratos, Jessie Huang, James N. Palmer, Nithin D. Adappa, Michael A. Kohanski, Darrell N. Kotton, Robert H Silverman, Wenli Yang, Edward Morrisey, Noam A. Cohen, Susan R Weiss

**Affiliations:** Departments of Microbiology, Perlman School of Medicine at the University of Pennsylvania, Philadelphia, PA, 19104; Medicine, Perlman School of Medicine at the University of Pennsylvania, Philadelphia, PA, 19104; Otorhinolaryngology, Perlman School of Medicine at the University of Pennsylvania, Philadelphia, PA, 19104; Department of Cancer Biology, Lerner Research Institute, Cleveland Clinic, Cleveland, OH 44195 USA; Department of Medicine, the Pulmonary Center, Center for Regenerative Medicine Boston University School of Medicine, Boston, MA 02118, USA; Corporal Michael J. Crescenz VA Medical Center, Philadelphia PA, 19104; Monell Chemical Senses Center, Philadelphia, PA, 19104; Institute for Regenerative Medicine, Perlman School of Medicine at the University of Pennsylvania, Philadelphia, PA, 19104; Penn-CHOP Lung Biology Institute, Perlman School of Medicine at the University of Pennsylvania, Philadelphia, PA, 19104; Penn Center for Research on Coronaviruses and Other Emerging Pathogens, Perlman School of Medicine at the University of Pennsylvania, Philadelphia, PA, 19104

**Keywords:** SARS-CoV-2, interferon, interferon signaling genes, OAS-RNase L, PKR

## Abstract

Coronaviruses are adept at evading host antiviral pathways induced by viral double-stranded RNA, including interferon (IFN) signaling, oligoadenylate synthetase–ribonuclease L (OAS-RNase L), and protein kinase R (PKR). While dysregulated or inadequate IFN responses have been associated with severe coronavirus infection, the extent to which the recently emerged SARS-CoV-2 activates or antagonizes these pathways is relatively unknown. We found that SARS-CoV-2 infects patient-derived nasal epithelial cells, present at the initial site of infection, induced pluripotent stem cell-derived alveolar type 2 cells (iAT2), the major cell type infected in the lung, and cardiomyocytes (iCM), consistent with cardiovascular consequences of COVID-19 disease. Robust activation of IFN or OAS-RNase L is not observed in these cell types, while PKR activation is evident in iAT2 and iCM. In SARS-CoV-2 infected Calu-3 and A549^ACE2^ lung-derived cell lines, IFN induction remains relatively weak; however activation of OAS-RNase L and PKR is observed. This is in contrast to MERS-CoV, which effectively inhibits IFN signaling as well as OAS-RNase L and PKR pathways, but similar to mutant MERS-CoV lacking innate immune antagonists. Remarkably, both OAS-RNase L and PKR are activated in *MAVS* knockout A549^ACE2^ cells, demonstrating that SARS-CoV-2 can induce these host antiviral pathways despite minimal IFN production. Moreover, increased replication and cytopathic effect in *RNASEL* knockout A549^ACE2^ cells implicates OAS-RNase L in restricting SARS-CoV-2. Finally, while SARS-CoV-2 fails to antagonize these host defense pathways, which contrasts with other coronaviruses, the IFN signaling response is generally weak. These host-virus interactions may contribute to the unique pathogenesis of SARS-CoV-2.

**Significance:** SARS-CoV-2 emergence in late 2019 led to the COVID-19 pandemic that has had devastating effects on human health and the economy. Early innate immune responses are essential for protection against virus invasion. While inadequate innate immune responses are associated with severe COVID-19 diseases, understanding of the interaction of SARS-CoV-2 with host antiviral pathways is minimal. We have characterized the innate immune response to SARS-CoV-2 infections in relevant respiratory tract derived cells and cardiomyocytes and found that SARS-CoV-2 activates two antiviral pathways, oligoadenylate synthetase–ribonuclease L (OAS-RNase L), and protein kinase R (PKR), while inducing minimal levels of interferon. This in contrast to MERS-CoV which inhibits all three pathways. Activation of these pathways may contribute to the distinctive pathogenesis of SARS-CoV-2.

## Introduction

Severe acute respiratory syndrome coronavirus (SARS-CoV)-2 emerged in China in late 2019, causing the COVID-19 pandemic with extensive morbidity and mortality, leading to major changes in day-to-day life in many parts of the world. This was the third lethal respiratory human coronavirus, after SARS-CoV in 2002 and Middle East respiratory syndrome coronavirus (MERS-CoV) in 2012, to emerge from bats in the twenty-first century. Although these viruses are all members of the *Betacoronavirus* genus (1), each has caused a somewhat different pattern of pathogenesis and spread in humans, with SARS-CoV-2 alone capable of spreading from asymptomatic or presymptomatic individuals (2). Therefore it is important to understand how these viruses interact with their host.

Coronaviruses are enveloped viruses with large, positive-sense single-stranded (ss)RNA genomes of around 30kb that can infect a diverse range of mammals and other species. Coronaviruses use much of their genomes, including their approximately 20 kb Orf1ab replicase locus comprising the 5’ two thirds of the genome, to encode proteins that antagonize host cell responses (3). As a result they are remarkably adept at antagonizing host responses, in particular the double-stranded RNA (dsRNA)-induced pathways that are essential components of the host innate immune response (4–8). In addition, interspersed among the structural genes encoded in the 3’ third of the genome are lineage-specific genes encoding accessory proteins, which are non-essential for RNA replication and variable among CoV lineages that further divide the *Betacoronavirus* genus (9). These accessory proteins often have functions in antagonizing host cell responses and thus likely contribute to discrepancies in pathogenesis and tropism observed among the different lineages (10–12).

Like other RNA viruses, coronaviruses produce dsRNA early during the infection cycle as a result of genome replication and mRNA transcription (13). Host cell pattern recognition receptors (PRRs) sense viral dsRNA as pathogenic non-self and respond by activating several antiviral pathways critical for early defense against viral invasion. DsRNA sensing by cytosolic PRRs can be divided into three key pathways – interferon (IFN) production, oligoadenylate-ribonuclease L (OAS-RNase L) activation, and protein kinase R (PKR) activation (**Fig 1**) (14). Detection of dsRNA by MDA5 during coronavirus infection (15), leads to the production of type I (α/β) and type III (λ) IFN. Upon binding to its specific cell surface receptor, IFN triggers phosphorylation of STAT1 and STAT2 transcription factors, which then induce expression of IFN stimulated genes (ISGs) with antiviral activities (16, 17). In parallel, dsRNA is also sensed by oligoadenylate synthetases (OASs), primarily OAS3, which synthesize 2’,5’-linked oligoadenylates (2-5A) (18, 19). Generation of 2-5A induces dimerization and activation of RNase L, leading to degradation of viral and host ssRNA (20). Finally, dsRNA sensing by PKR induces PKR autophosphorylation, permitting PKR to then phosphorylate the translation initiation factor eIF2α, which results in protein synthesis shutdown and restriction of viral replication (21). While RNase L and PKR antiviral activity are not dependent on IFN production (18), the genes encoding OASs and PKR are ISGs, therefore these pathways can be activated and/or reinforced by IFN production. Similarly, RNase L and PKR activation can promote IFN production, cellular stress, inflammation, and/or apoptotic death (22–27), thus further reducing host cell viability.

**Figure 1.**
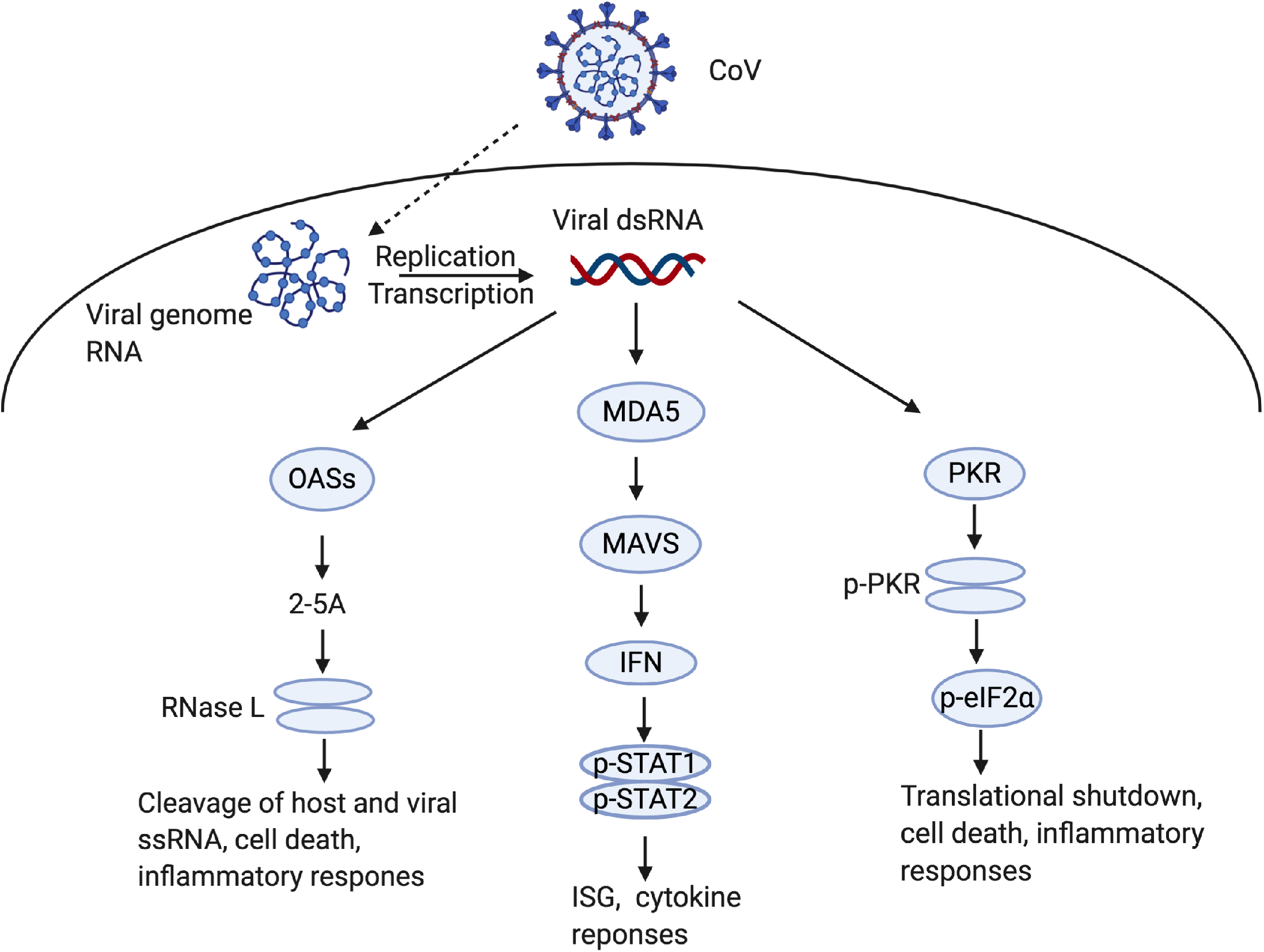
Double-stranded RNA induced innate immune responses during SARS-CoV-2 infection. Coronavirus double-stranded RNA (dsRNA) is produced through replication and transcription and recognized by cytosolic OAS, MDA5, or PKR host receptors to activate innate immune pathways. MDA5 signals through MAVS, leading to type I and type III IFN production and release from the cell where it binds to cell surface receptors, which induces phosphorylation and heterodimerization of STAT1 and STAT2 that then prompt ISG transcription and cytokine responses. OASs produce 2’-5’-oligoadenylates (2-5A) that bind RNase L, leading to homodimerization and catalytic activation of RNase L, which cleaves host and viral ssRNA to trigger apoptosis and inflammation. PKR autophosphorylates before phosphorylating eIF2α, which leads to translational arrest, cell death, and inflammatory responses. Graphic was created with Biorender.com

Induction and inhibition of innate immune responses during infection with SARS-CoV-2 have yet to be fully characterized. Several recent reports implicate genetic deficiencies in IFN responses (28, 29) or polymorphisms in OAS genes (30) with more severe COVID-19 disease, emphasizing the importance of understanding the interactions between SARS-CoV-2 and these innate response pathways. Furthermore, while it is known that SARS-CoV-2 enters the human body through the upper respiratory tract, it is unclear which cell types of the upper and lower respiratory system contribute to sustained infection and resulting disease in the airways and elsewhere. We have performed SARS-CoV-2 infections of primary nasal epithelial cells, induced pluripotent stem cell (iPSC)-derived alveolar type 2 cells (iAT2), and iPSC-derived cardiomyocytes (iCM), which collectively represent the host tissues likely affected by clinical SARS-CoV-2 infection (31, 32). We assessed viral replication in these cell types as well as the degree of ensuing dsRNA-sensing responses. We also employed two lung derived immune-competent cells lines, Calu-3 and A549 cells, to investigate dsRNA-induced pathway activation during SARS-CoV-2 infection. In addition, we compared host responses to SARS-CoV-2 with those of MERS-CoV and MERS-CoV-ΔNS4ab, a mutant lacking expression of two dsRNA-induced innate immune pathway antagonists that we have characterized previously (10).

## Results

### SARS-CoV-2 replicates efficiently in cells derived from upper and lower respiratory tract

We compared the replication of SARS-CoV-2 and MERS-CoV in nasal epithelia-derived cells, a relevant site of infection *in vivo* (**Fig 2A**). For each virus, replication was similar in cells from four different individuals, although the extent of replication was somewhat variable. The trends in replication kinetics, however, were significantly different between SARS-CoV-2 and MERS-CoV infections. Replication of SARS-CoV-2 increased until 96hpi, but then plateaued at nearly 10^6^ plaque-forming units (PFU)/ml. MERS-CoV replication peaked at 96hpi, at a lower titer than SARS-CoV-2, and produced fewer PFU/mL at later timepoints. Nasal epithelial cell cultures were stained with antibodies to identify ciliated cells (anti-type IV β-tubulin), a key feature of this cell type, and either SARS-CoV-2 or MERS-CoV nucleocapsid expression (anti-N protein) (**Fig 2B**). We detected abundant N expression in both SARS-CoV-2 and MERS-CoV infected cells, indicating that these cells were sufficiently infected at 48 hours post infection (hpi). Interestingly, robust replication occurred in these cultures, despite a very low level of ACE2 protein expression in cells from the three individuals examined (**Fig 2C**).

**Figure 2.**
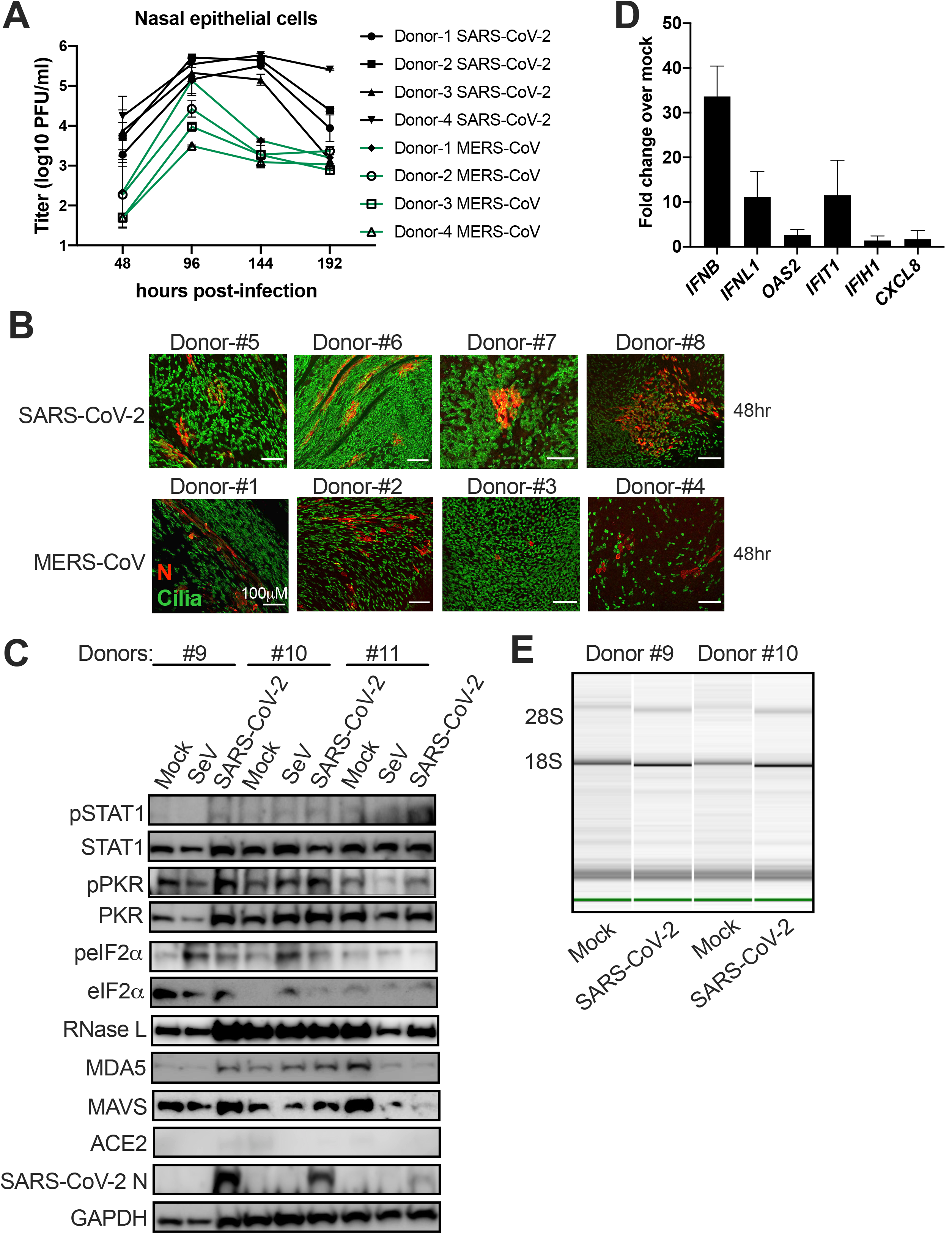
Infection of nasal epithelia-derived cells by SARS-CoV-2 and MERS-CoV. Nasal cells were cultured in air-liquid trans-wells, and mock infected or infected with SARS-CoV-2 (MOI=5), MERS-CoV (MOI=5), or Sendai Virus (SeV), MOI=10, apically. (A) At indicated times, apically released virus was quantified by plaque assay on Vero-E6 cells. Values are means ± SD (error bars). Statistical significance (not displayed) was determined by two-way ANOVA (*, P < 0.05). One experiment was performed using four separate donors. (B) At 48 hpi, nasal cells were fixed with 4% PFA and permeabilized. Expression of nucleocapsid (N) protein (red) of SARS-CoV-2 and MERS-CoV was detected with an anti-N antibody, and cilia (green) with an anti-type IV β-tubulin antibody by immunofluorescence assay (IFA). One representative image is shown from at least three independent experiments, with four donors for each virus infection shown. Scale bar = 100μm. (C) At 120 hpi, cells were lysed, and proteins were analyzed by immunoblotting with antibodies as indicated. One experiment using three separate donors was performed. (D) At 120 hpi, total RNA was harvested, and the mRNA expression level of *IFNB, IFNL1, OAS2, IFIT1, IFIH1, CXCL8*was quantified by RT-qPCR. Cycle threshold (C_T_) values were normalized to 18S rRNA to generate ΔC_T_ values (ΔC_T_ = C_T_ gene of interest - C_T_ 18S rRNA). Fold change over mock values were calculated by subtracting mock infected ΔCT values from virus infected ΔC_T_ values, displayed as 2^-Δ(ΔCt)^. Technical replicates were averaged, the means for each replicate displayed, ± SD (error bars). One experiment was performed using three separate donor samples. (E) Total RNA was harvested from two donors at 120 hpi and rRNA integrity determined by Bioanalyzer. The position of 28S and 18S rRNA and indicated. Data shown are from one representative experiment of two independent experiments. (See also Figures S1A&S2).

We measured dsRNA-induced host responses to SARS-CoV-2 infection, including type I and type III IFN mRNA induction, RNase L activation, and PKR activation, in the nasal cells. For RT-qPCR analysis, we extracted RNA from SARS-CoV-2 infected cultures from four different donors at 120hpi. We verified that virus was replicating by quantifying viral genome copies from intracellular RNA (**Fig S1A**). We then quantified mRNA expression of IFN-β (type I IFN), IFN-λ (type III IFN), select ISGs (*OAS2, IFIT1, IFIH1*), and the neutrophil attracting chemokine IL-8 (*CXCL8*), which has been implicated in nasal inflammation during viral infection (33, 34) (**Fig 2D**). There was some induction of IFN-β and to a lesser extent IFN-λ mRNA, and minimal induction of the ISG mRNAs examined. Similarly, *CXCL8* encoding IL-8 was barely induced. Interestingly, this may be at least partially due to high basal levels of IFN (notably IFN-λ) and ISG (notably OAS2) mRNAs compared with other cell types examined below, which would result in weak fold changes in mRNA levels compared with mock infected cells (**Fig S2**). To further investigate this very weak ISG induction, using cells from the same donors as the IFN/ISG mRNA quantification, we assessed the phosphorylation of STAT1, a transcription factor that is itself encoded by an ISG, which is primarily a key mediator of type I and type III IFN signaling (35). Consistent with the weak activation of ISGs, there was no evidence of phosphorylation of STAT1 (**Fig 2C**). In addition, we did not detect PKR activation in SARS-CoV-2 infected cells, as indicated by the absence of phosphorylated PKR and eIF2α. This is in contrast to the phosphorylated eIF2α detected in Sendai virus (SeV) infected cells from two of the three donors (**Fig 2C**). We also assessed activation of the OAS-RNase L pathway during SARS-CoV-2 infection of cells from two of the same four donors. Since 28S and 18S ribosomal RNAs (rRNAs) are targeted for degradation by activated RNase L, we evaluated 28S and 18S rRNA integrity using a Bioanalyzer as a readout for RNase L activation. The absence of any rRNA degradation in SARS-CoV-2 infected cells (**Fig 2E**) indicated that RNase L was not activated despite abundant RNase L protein expression (**Fig 2C**).

Next, we sought to examine host innate immune responses during infection of alveolar type 2 cells (AT2), a major target of SARS-CoV-2 infection in humans (31, 36, 37). We employed induced pluripotent stem cell (iPSC)-derived iAT2 cells (SPC2 line), expressing tdTomato from the endogenous locus of surfactant protein-C (SFTPC), an AT2 cell specific marker (38). As in nasal cells, virus replicated efficiently, reaching a titer of 10^6^ PFU/ml by 48hpi (**Fig 3A**). Staining of cultures with an anti-N antibody showed that most of the iAT2 cells were infected, without obvious cytopathic effect (CPE) during infection (**Fig 3B**). Notably, SARS-CoV-2 infection of iAT2 cells was robust despite ACE2 expression being below the level of detection by immunoblotting (**Fig 3C**). We observed activation of the PKR pathway as indicated by both PKR and eIF2α phosphorylation (**Fig 3C**). We extracted RNA from infected iAT2 cells for RT-qPCR analysis, verified these cells were replicating virus by quantifying genome RNA copies (**Fig S1B**), and assessed IFN/ISG induction. As with the nasal cells, we observed weak induction of IFN-β and IFN-l mRNA from mock infected and infected cells (**Fig 3D**), as well as no detection of MDA5 protein (15), a dsRNA sensor in the pathway leading to IFN production during coronavirus infection (**Fig 3C)**. We used the alphavirus Sindbis virus (SINV) as a positive control, which we have previously shown induces robust activation of all dsRNA-induced pathways (10). Surprisingly, we observed greater increases in OAS2 and IFIT mRNA expression by SARS-CoV-2 compared with SINV (**Fig 3D**), but with minimal induction of IFIH1 mRNA, consistent with the lack of MDA5 protein expression (**Fig 3C&D)**. However, we did not observe phosphorylation of STAT1 (**Fig 3C**), as in the nasal cells above. Additionally, we did not observe any degradation of rRNA in SARS-CoV-2 infected cells, and only slight degradation by SINV despite ample expression of RNase L (**Fig 3E**), suggesting minimal activation of RNase L in iAT2 cells in general.

**Figure 3.**
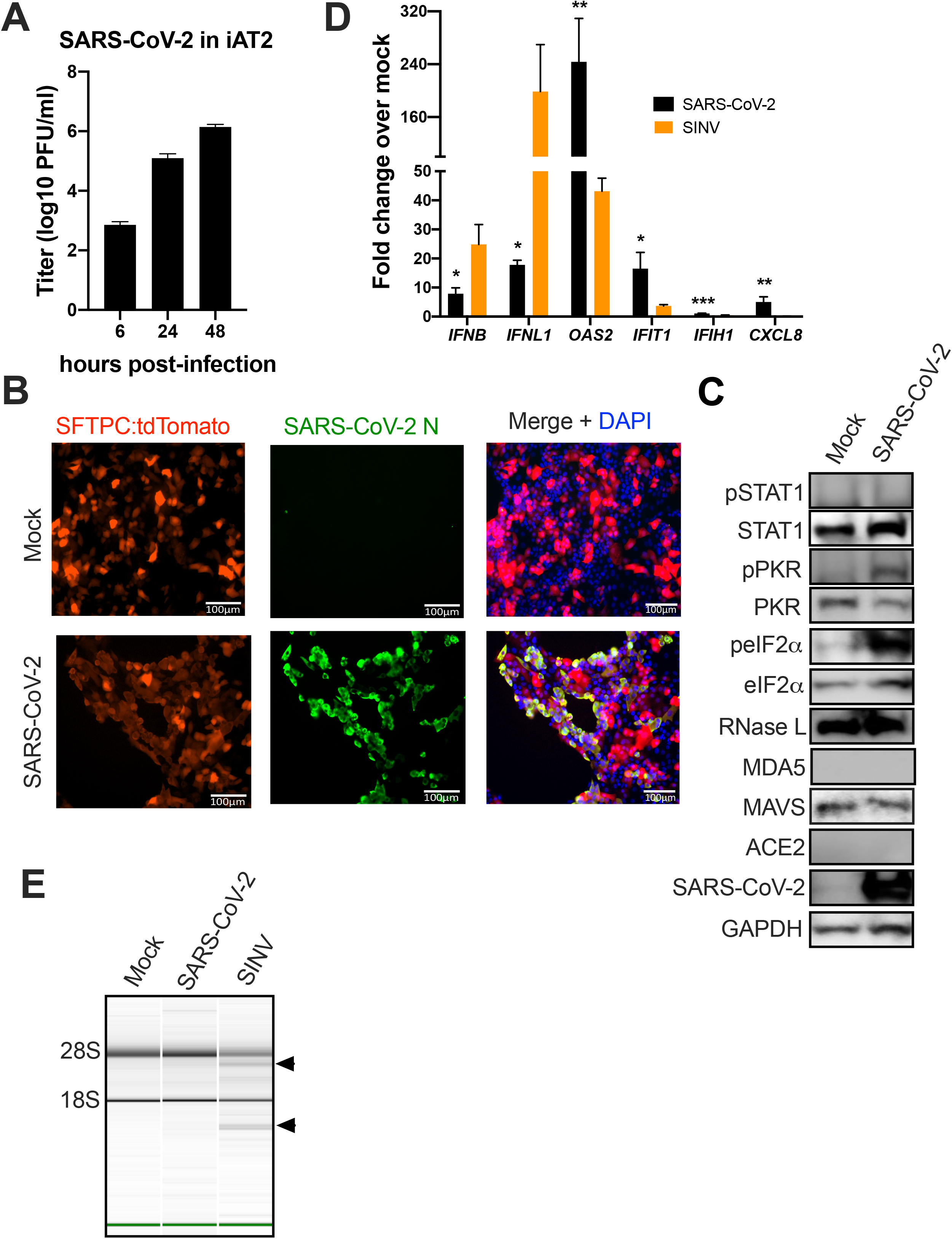
Infection of iPSC-derived AT2 cells (iAT2) by SARS-CoV-2. iAT2 cells were mock infected or infected with SARS-CoV-2 at MOI=5 or SINV at MOI=1. (A) At indicated times, supernatants were collected and infectious virus was quantified by plaque assay on Vero-E6 cells. Values are means ± SD (error bars). Data shown are one representative experiment from at least three independent experiments. (B) At 48 hpi, cells were fixed with 4% PFA and permeabilized. Expression of nucleocapsid (N) protein (green) of SARS-CoV-2 and the expression of SFTPC promoter control tdTomato fluorescent protein (AT2 marker in red) was examined by IFA. Channels are merged with DAPI nuclear staining. Images shown are representative from at least three independent experiments. Scale bar = 100μm. (C) At 48 hours post infection, cells were lysed and proteins were analyzed by immunoblotting with antibodies as indicated. Data shown are from one representative experiment of two independent experiments. (D) At 16 (SINV) or 48 (SARS-CoV-2) hpi, total RNA was harvested, and the mRNA expression level of *IFNB, IFNL1, OAS2, IFIT1, IFIH1, CXCL8* was quantified by RT-qPCR. C_T_ values were normalized to 18S rRNA to generate ΔC_T_ values (ΔC_T_ = C_T_ gene of interest - C_T_ 18S rRNA). Fold change over mock values were calculated by subtracting mock infected ΔC_T_ values from virus infected ΔC_T_ values, displayed as 2^-Δ(ΔCt)^. Technical replicates were averaged, the means for each replicate displayed, ± SD (error bars). Statistical significance was determined by Student *t* test (*, P < 0.05; **, P < 0.01; ***, P < 0.001). Data shown are from one representative experiment of two independent experiments. (E) Total RNA was harvested at 16 (SINV) or 48 (SARS-CoV-2) hpi and rRNA integrity determined by Bioanalyzer. The position of 28S and 18S rRNA and indicated. Data shown are from one representative experiment of two independent experiments. (See also Figures S1B&S2).

### SARS-CoV-2 replicates and induces innate immune responses in iPSC-derived cardiomyocytes

Since many COVID-19 patients experience cardiovascular symptoms and pathology (39, 40), we investigated SARS-CoV-2 infection of iPSC derived-cardiomyocytes (iCM). SARS-CoV-2 replicated robustly in these cells, reaching titers of approximately 10^6^ PFU/ml by 48hpi (**Fig 4A**), similar to replication in nasal and iAT2 cells. Cells were stained with an antibody against cardiac troponin-T (cTnT) as a marker for cardiomyocytes, and an antibody against the viral N protein to identify infected cells (**Fig 4B**). In addition, we detected clear CPE in the iCM, which differed from infected nasal and iAT2 cells. This CPE included syncytia resulting from cell-to-cell fusion, which is typical of coronaviruses (41–45). Interestingly, while we observed detectable ACE2 protein expression in mock infected or SINV infected cells in two independent experiments, we observed loss of ACE2 expression upon SARS-CoV-2 infection, consistent with a recent study (32) (**Fig 4C**). We extracted RNA from mock infected cells and cells infected with SARS-CoV-2 or SINV, verified that virus was replicating by quantifying viral genome (**Fig S1C**), and quantified expression of mRNAs for IFNs and select ISGs. We found low levels of IFN/ISGs transcript in iCM similar to the nasal and iAT2 cells (**Fig D**), perhaps due to the undetectable levels of MDA5 and MAVS protein expression in these cells (**Fig 4C**). SINV also induced host mRNAs weakly, with the exception of IFN-λ, in these cells (**Fig 4D**). We observed no degradation of rRNA, suggesting an absence of RNase L activation in iCM with SARS-CoV-2 or SINV (**Fig 4E**), despite clear infection with either virus (**Fig S1C**). This was not surprising as there was no RNase L detectable by immunoblot in these cells (**Fig 4C**). Finally, as in iAT2 cells, we observed phosphorylation of PKR and eIF2α, indicating that the PKR antiviral pathway is activated (**Fig 4C**).

**Figure 4.**
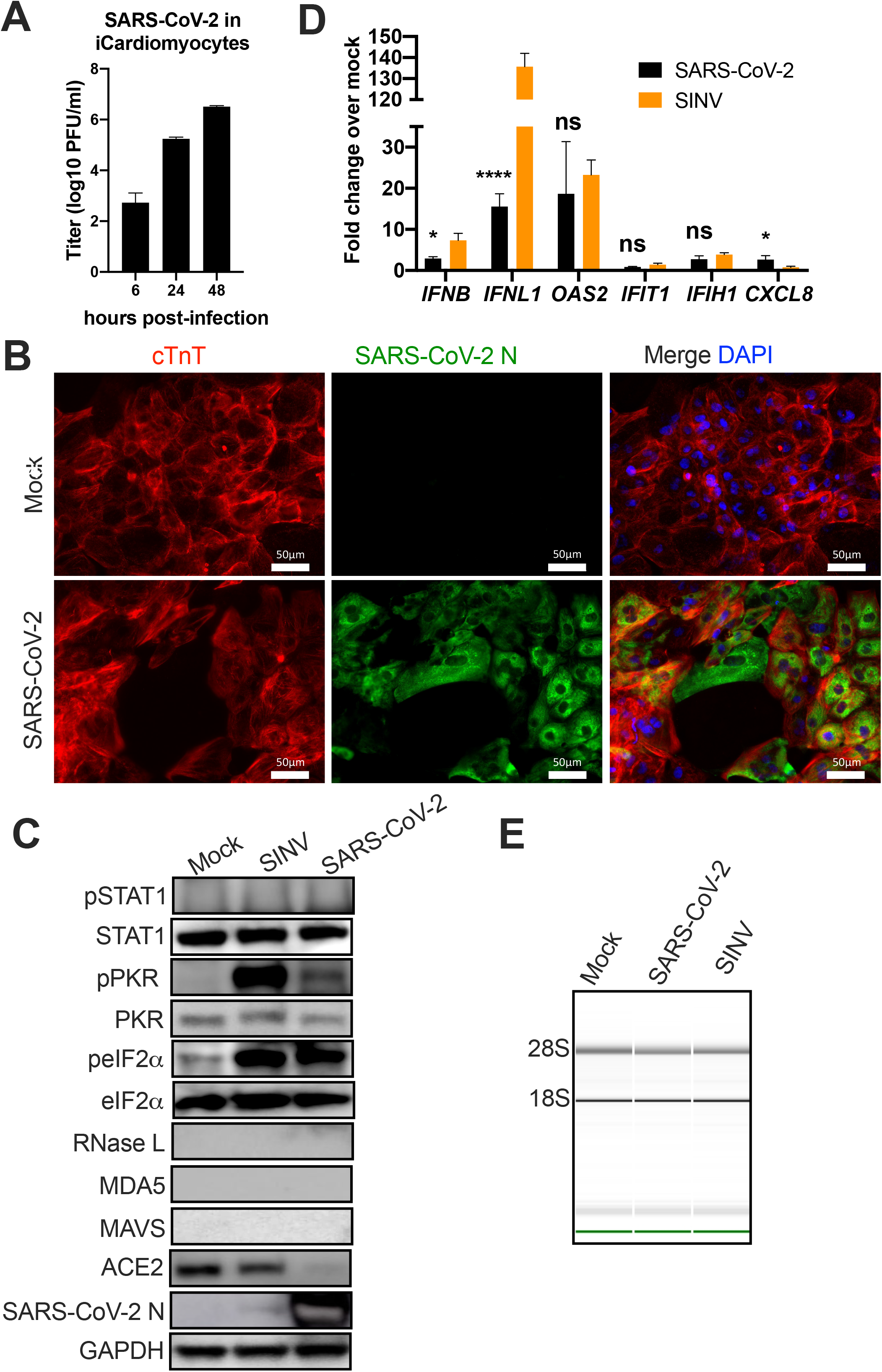
Infection of iPSC-derived cardiomyocytes (iCM) by SARS-CoV-2. iCM were mock infected or infected at MOI=1 with SARS-CoV-2 or SINV. (A) At indicated times, supernatants were collected and virus quantified by plaque assay on Vero-E6 cells. Values are means ± SD (error bars). Data shown are one representative experiment from at least three independent experiments. (B) At 48 hpi, iCM were fixed with 4% PFA and permeabilized, the expression of SARS-CoV-2 N (green) of and of cTnT protein (cardiomyocyte marker, red) was examined by IFA. Channels are merged with DAPI nuclear staining. Images shown are representative from three independent experiments. Scale bar = 50μm. (C) At 16 (SINV) or 48 (SARS-CoV-2) hpi, cells were lysed and proteins were analyzed by immunoblotting with antibodies as indicated. Immunoblots were performed at least two times and one representative blot is shown. (D) At 16 (SINV) or 48 (SARS-CoV-2) hpi, total RNA was harvested, the mRNA expression level of *IFNB, IFNL1, OAS2, IFIT1, IFIH1, CXCL8* was quantified by RT-qPCR. C_T_ values were normalized to 18S rRNA to generate ΔC_T_ values (ΔC_T_ = C_T_ gene of interest - C_T_ 18S rRNA). Fold change over mock values were calculated by subtracting mock infected ΔC_T_ values from virus infected ΔC_T_ values, displayed as 2^-Δ(ΔCt)^. Technical replicates were averaged, the means for each replicate displayed, ± SD (error bars). Statistical significance was determined by Student *t* test (*, P < 0.05; ****, P < 0.0001; ns = not significant). Data shown are from one representative experiment of two independent experiments. (E) Total RNA was harvested at 16 (SINV) or 48 (SARS-CoV-2) hpi, and rRNA integrity determined by Bioanalyzer. The position of 28S and 18S rRNA and indicated. Data shown are from one representative experiment of two independent experiments. (See also Figures S1C&S2).

### SARS-CoV-2 replicates in respiratory epithelial cell lines and induces dsRNA responsive pathways

To further characterize the relationship between SARS-CoV-2 and dsRNA-induced host response pathways, we chose two respiratory epithelium-derived human cell lines, A549 and Calu-3, both of which are immune competent and have been used for studies of SARS-CoV (46) and MERS-CoV (10, 47). A549 cells were not permissive to SARS-CoV-2, due to lack of expression of the SARS-CoV-2 receptor ACE2 (**Fig S3**). Therefore, we generated A549 cells expressing the ACE2 receptor (A549^ACE2^) by lentiviral transduction, and used two single cell clones, C44 and C34, for all experiments (**Fig S3**). Both A549^ACE2^ clones express high levels of ACE2 greater than the endogenously expressed ACE2 in Calu-3 cells (**Fig S3**) and in the primary cells discussed above (**Fig 2–4**).

We performed single step growth curves to measure replication of SARS-CoV-2 over the course of one infectious cycle in A549^ACE2^ cells, simian Vero-E6 cells, which are commonly used to prepare SARS-CoV-2 stocks, and Calu-3 cells (clone HTB-55). SARS-CoV-2 replicated robustly in A549^ACE2^ and Vero-E6 cells (**Fig 5A**), although viral yields were lower in Calu-3 cells (**Fig 5B**). Since Calu-3 cells also support MERS-CoV infection, we compared SARS-CoV-2 replication to that of wild type MERS-CoV and MERS-CoV-ΔNS4ab, a mutant deleted in host cell antagonists NS4a, a dsRNA-binding protein, and NS4b, a 2’5’-phosphodiesterase that prevents RNase L activation and nuclear translocation of NF-κB (10, 48). Consistent with our previous work (10), MERS-CoV-ΔNS4ab reduced viral titers from WT MERS-CoV levels, although they remained higher than SARS-CoV-2 titers (**Fig 5B**). To further understand the replication of SARS-CoV-2, we stained A549, Vero-E6, and Calu-3 cells at 24 hpi with antibodies against viral N protein and viral dsRNA, including additional Calu-3 staining at 48 hpi since replication kinetics are slower (**Fig 5C**). We observed cytopathic effect in all three cell types, with N localized to the cytoplasm. Syncytia were observed in A549^ACE2^ and Calu-3 cells, but not in Vero-E6 cells (**Fig 5C**). We also observed viral dsRNA localized to perinuclear foci as we and others have described during infection with other coronaviruses (10, 49–51).

**Figure 5.**
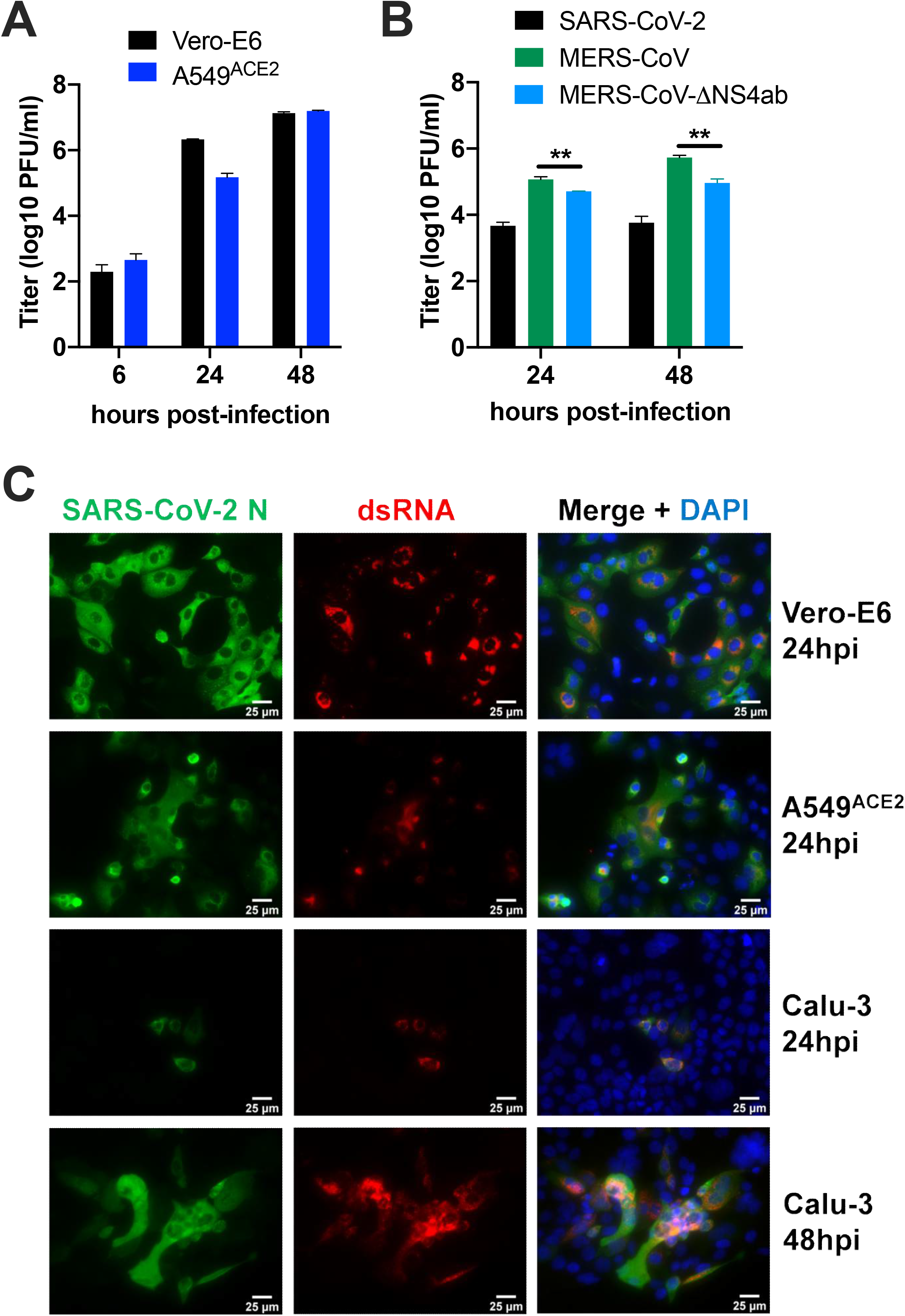
Replication of SARS-CoV-2 in A549^ACE2^ and Calu-3 cell lines. (A) Vero-E6 or A549^ACE2^ (clone 44) cells were infected with SARS-CoV-2 at MOI=1. At the indicated times, supernatant was collected and virus quantified by plaque assay on Vero-E6 cells. Values are means ± SD (error bars). (B) Calu-3 cells were infected with SARS-CoV-2, MERS-CoV or MERS-CoV-ΔNS4ab at MOI=1. Supernatant was collected at the indicated times and virus quantified by plaque assay on Vero-E6 cells (SARS-CoV-2) or VeroCCL81 cells (MERS-CoV and MERS-CoV-Δ4ab). Values represent means ± SEM (error bars). Statistical significance was determined by Student *t* test (**, P < 0.01). Data shown are one representative experiment of three independent experiments. (C) Vero-E6, A549^ACE2^ (clone 34), and Calu-3 cells were grown on untreated (Vero-E6 and A549^ACE2^) or collagen-coated (Calu-3) glass coverslips before infection with SARS-CoV-2 at MOI = 1. At indicated hpi, cells were fixed with 4% PFA and permeabilized for N (green) and dsRNA (red) expression detection by IFA using anti-N and J2 antibodies, respectively. Channels are merged with DAPI nuclear staining. Images shown are representative from two independent experiments. Scale bar = 25μm. (See also Figure S3&S4A).

We used RT-qPCR to quantify the induction of type I and type III IFNs and select ISGs at 24 and 48 hpi (**Fig 6A**), as well as the intracellular viral genome copies to verify replication (**Fig 6B**) in A549^ACE2^ cells. Using SINV as a positive control, we found relatively low levels of both IFNβ and IFNλ mRNA at 24 and 48 hpi by SARS-CoV-2, compared to SINV (**Fig 6A**). Notably, IFN induction was greater than observed in the nasal, iAT2, or iCM cells, possibly due to lower basal levels of IFNβ, but not IFNλ, mRNA in the A549^ACE2^ cells, which allow for greater fold changes over mock infected cells (**Fig S2**). Levels of ISG mRNAs were variable, with SARS-CoV-2 inducing moderate levels of OAS2 and IFIT1 mRNAs, but only late in infection (48 hpi), similar to those induced by SINV at 24 hpi **(Fig 6A**). We observed minimal effects on mRNA levels of IFIH1 and CXCL8 at both timepoints (**Fig 6A**). Furthermore, we did not detect any STAT1 phosphorylation at 24 hpi (**Fig 6C**), which correlates with weak ISG expression, suggesting defective IFN signaling downstream of IFN production.

**Figure 6.**
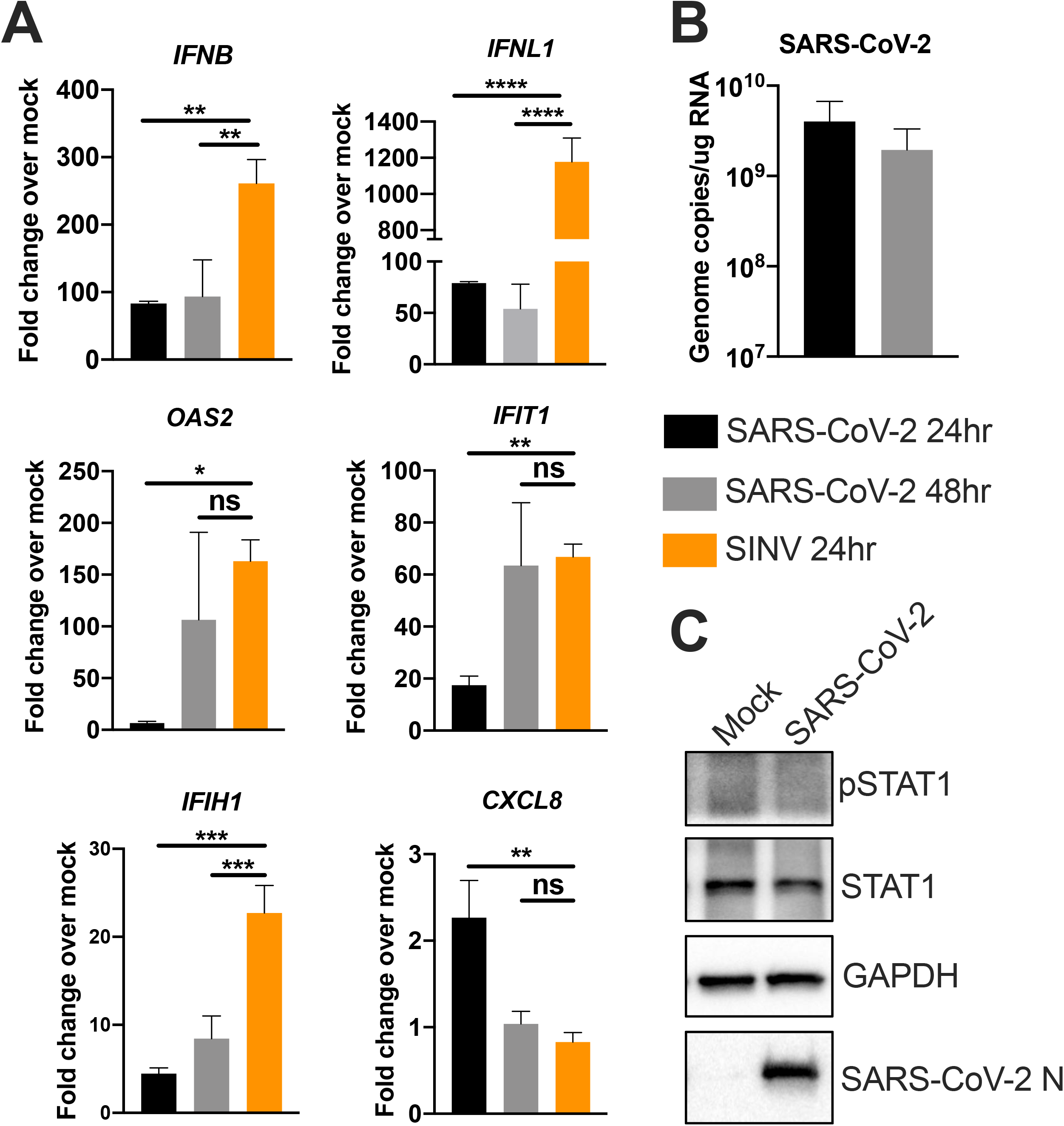
SARS-CoV-2 IFN responses in the lung epithelia-derived A549^ACE2^ cell line. A549^ACE2^ cells were mock infected or infected with SINV (MOI=1) or SARS-CoV-2 (MOI=5). (A) Total RNA was harvested at 24 and 48 hpi. Expression of IFNB, IFNL1, OAS2, IFIT1, IFIH1, and CXCL8 mRNA was quantified by RT-qPCR. C_T_ values were normalized to 18S rRNA to generate ΔC_T_ values (ΔC_T_ = C_T_ gene of interest - C_T_ 18S rRNA). Fold change over mock values were calculated by subtracting mock infected ΔC_T_ values from virus infected ΔC_T_ values, displayed as 2^-Δ(ΔCt)^. Technical replicates were averaged, the means for each replicate displayed, ± SD (error bars). (B) Viral genome copies per ug of total RNA were calculated at 24 and 48hpi by RT-qPCR standard curve generated using a digested plasmid encoding SARS-CoV-2 nsp12. Values are means ± SD (error bars). Statistical significance was determined by one-way ANOVA (*, P < 0.05; **, P < 0.01; ***, P < 0.001; ****, P < 0.0001; ns = not significant). (C) At 24 hpi, A549^ACE2^ cells were lysed and proteins harvested. Protein expression was analyzed by immunoblot using the indicated antibodies. All data are one representative experiment of three independent experiments, carried out with A549^ACE2^ clone 44. (See also Figures S2, S4B&C).

We evaluated IFN/ISG responses in Calu-3 cells, which provided a second lung-derived cell line that additionally supports both SARS-CoV-2 and MERS-CoV infection, allowing us to compare host responses between the two lethal CoVs. We compared SARS-CoV-2 responses to both WT MERS-CoV and mutant MERS-CoV-ΔNS4ab (**Fig 7A**). Although we observed reduced MERS-CoV-ΔNS4ab infectious virus production compared with WT MERS-CoV (**Fig 5B**), we detected similar intracellular viral genome levels of all three viruses (**Fig 7B**). We found previously that MERS-CoV-ΔNS4ab induces higher levels of IFNs and ISGs compared to WT MERS-CoV, and also activates RNase L and PKR (10). Herein, in Calu-3 cells, we observed greater SARS-CoV-2 induction of IFN mRNAs as compared to A549^ACE2^ cells (**Fig 6A&S4B**). Interestingly, SARS-CoV-2 induced higher IFN mRNA levels than WT MERS-CoV at 24 and 48 hpi (**Fig 7A**). Similarly, SARS-CoV-2 generally induced more ISG mRNA than WT MERS-CoV, and even more OAS2 mRNA than MERS-ΔNS4ab (**Fig 7A**). Induction of CXCL8 was weak for all viruses (**Fig 7A**). Notably, SARS-CoV-2 induced ISG mRNAs in Calu-3 (24hpi) without the delay observed in A549^ACE2^ cells. Consistent with earlier ISG mRNA induction during infection, SARS-CoV-2 infection promoted phosphorylation of STAT1 in Calu-3 cells (**Fig 7C**), as recently reported (52). SARS-CoV-2 induced phosphorylation of STAT1 as well as rapid IFIT1 and OAS2 mRNA induction suggests a similar host response to SARS-CoV-2 as that observed during mutant MERS-CoV-ΔNS4ab infection, and not that of WT MERS-CoV infection.

**Figure 7.**
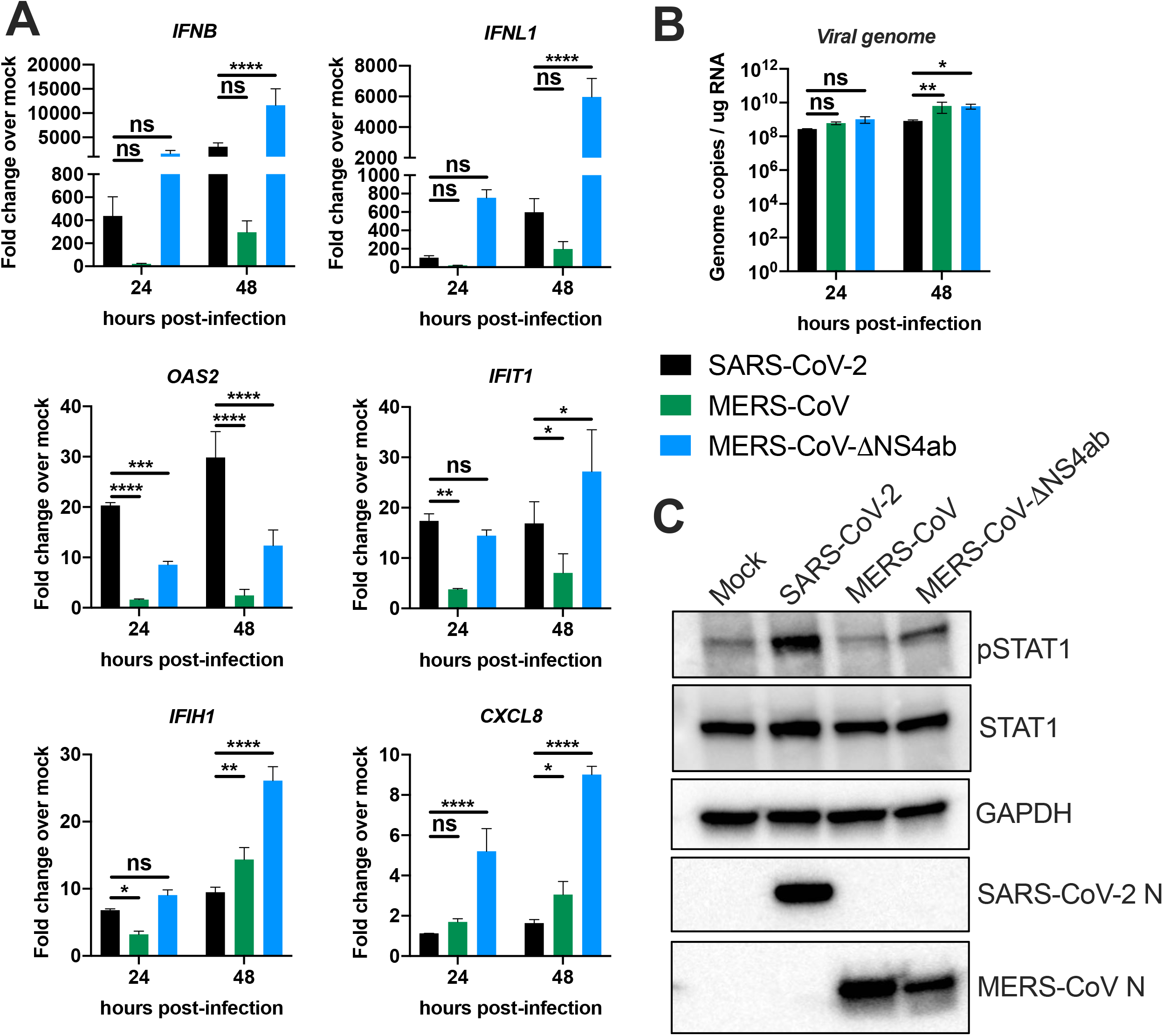
SARS-CoV-2 and MERS-CoV IFN responses in the lung-derived Calu-3 cells. Calu-3 cells were mock treated or infected with SARS-CoV-2, MERS-CoV or MERS-CoV-ΔNS4ab at MOI=5. (A) At 24 or 48 hpi, total RNA was harvested. Expression of *IFNB, IFNL1, OAS2, IFIT1, IFIH1*, and *CXCL8* mRNA was quantified by RT-qPCR. C_T_ values were normalized to 18S rRNA to generate ΔC_T_ values (ΔC_T_ = C_T_ gene of interest - C_T_ 18S rRNA). Fold change over mock values were calculated by subtracting mock infected ΔC_T_ values from virus infected ΔC_T_ values, displayed as 2^-Δ(ΔCt)^. Technical replicates were averaged, the means for each replicate displayed, ± SD (error bars). Statistical significance was determined by two-way ANOVA (*, P < 0.05; **, P < 0.01; ***, P < 0.001; ****, P < 0.0001; ns = not significant). (B) Viral genome copies per ug of total RNA were calculated by RT-qPCR standard curve generated using a digested plasmid encoding SARS-CoV-2 nsp12 or plasmid encoding a region of MERS-CoV orf1ab. Values are means ± SD (error bars). Statistical significance was determined by two-way ANOVA (*, P < 0.05; **, P < 0.01; ns = not significant). (C) At 24 hpi, Calu-3 cells were lysed and proteins harvested. Proteins were analyzed by immunoblotting using the indicated antibodies. All data are one representative experiment of three independent experiments. (See also Figure S2).

### SARS-CoV-2 infection activates RNase L and PKR

We assessed activation of the RNase L pathway by analyzing intracellular rRNA integrity in infected cells, as described above. We found that in A549^ACE2^, SARS-CoV-2 promoted rRNA degradation by 24 hpi, which was more clearly observed at 48 hpi, using SINV as a positive control (**Fig 8A**). Evaluation of RNase L activation in SARS-CoV-2, WT MERS-CoV, and MERS-CoV-ΔNS4ab infected Calu-3 cells showed SARS-CoV-2 activation of RNase L to a similar extent as MERS-CoV-ΔNS4ab (10, 53) (**Fig 8B**). In contrast, as we previously reported, MERS-CoV failed to activate RNase L (10, 47) (**Fig 8B**). We also observed activation of PKR as indicated by phosphorylation of PKR and downstream eIF2α, in both A549^ACE2^ cells (**Fig 8C**) and Calu-3 cells (**Fig 8D**) infected with SARS-CoV-2. In Calu-3 cells, SARS-CoV-2 induced PKR phosphorylation to a similar extent as MERS-CoV-ΔNS4ab, while WT MERS-CoV failed to induce a response. These data are consistent with IFN/ISG induction data described above, suggesting that SARS-CoV-2 may not antagonize dsRNA pathways as efficiently as MERS-CoV, but instead induces host responses similar to those observed during MERS-CoV-ΔNS4ab infection.

**Figure 8.**
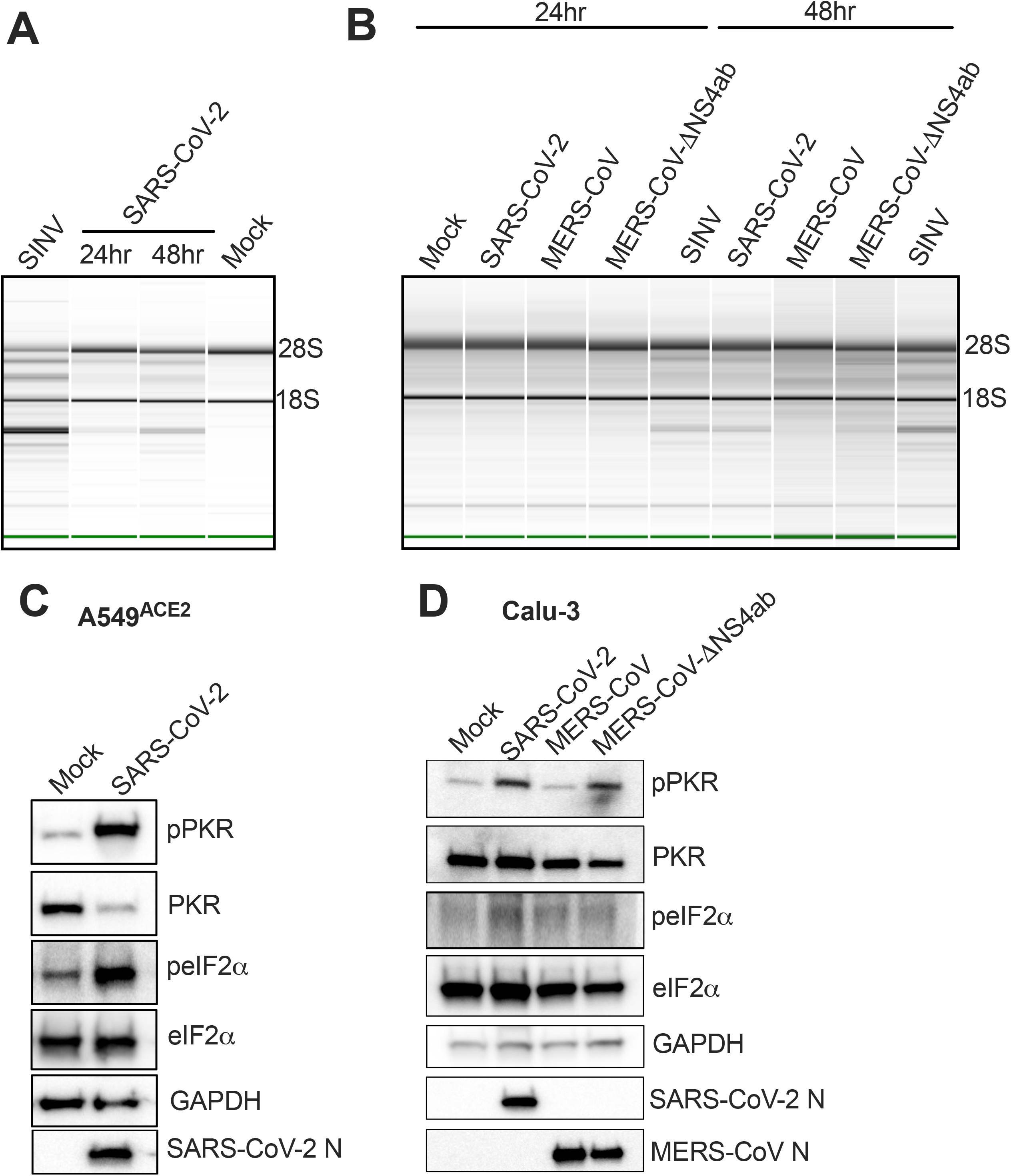
SARS-CoV-2 infection leads to activation of RNase L and PKR in A549^ACE2^ and Calu-3 cells. A549^ACE2^ and Calu-3 cells were mock infected or infected with SARS-CoV-2, MERS-CoV, or MERS-CoV-ΔNS4ab at MOI=5. Total RNA was harvested from A549^ACE2^ cells (A) or Calu-3 cells (B) at 24 and 48 hpi. 28S and 18S rRNA integrity was assessed by Bioanalyzer. 28S and 18s rRNA bands are indicated. At 24 hpi, A549^ACE2^ cells (C) or Calu-3 cells (D) were lysed and proteins harvested for analysis by immunoblotting using the indicated antibodies. All data are one representative experiment of three independent experiments. (See also Figure S4D&E).

The A549^ACE2^ cells were valuable in that they provided a system with intact innate immune responses that was also amenable to CRISPR-Cas9 engineering. Thus, we used the A549^ACE2^ cells to construct additional cell lines with targeted deletions of *MAVS, RNASEL*, or *PKR*, as we have done previously for parental A549 cells (19, 54). We could then use these cells to determine whether activation of IFN, RNase L, and/or PKR resulted in attenuation of SARS-CoV-2 replication (19, 54). We validated the knockout (KO) A549^ACE2^ cell lines by western blot (**Fig S5A**) and compared replication of SARS-CoV-2 in *MAVS* KO, *RNASEL* KO and *PKR* KO cells with levels in WT A549^ACE2^ cells (**Fig 9A**). Interestingly, there was little effect on SARS-CoV-2 replication with MAVS or PKR expression absent. At 48 hpi in *RNASEL* KO cells, virus replication was two-to four-fold higher compared to WT A549^ACE2^ cells **(Fig 9A**). While the difference in replication between *RNASEL* KO and WT was not extensive, it was statistically significant in three independent experiments. As a result of higher viral titers, infected *RNASEL* KO cells exhibited strikingly more CPE as compared with WT, *PKR* KO, or *MAVS* KO cells, as demonstrated by crystal violet-staining of infected cells (**Fig 9B**).

**Figure 9.**
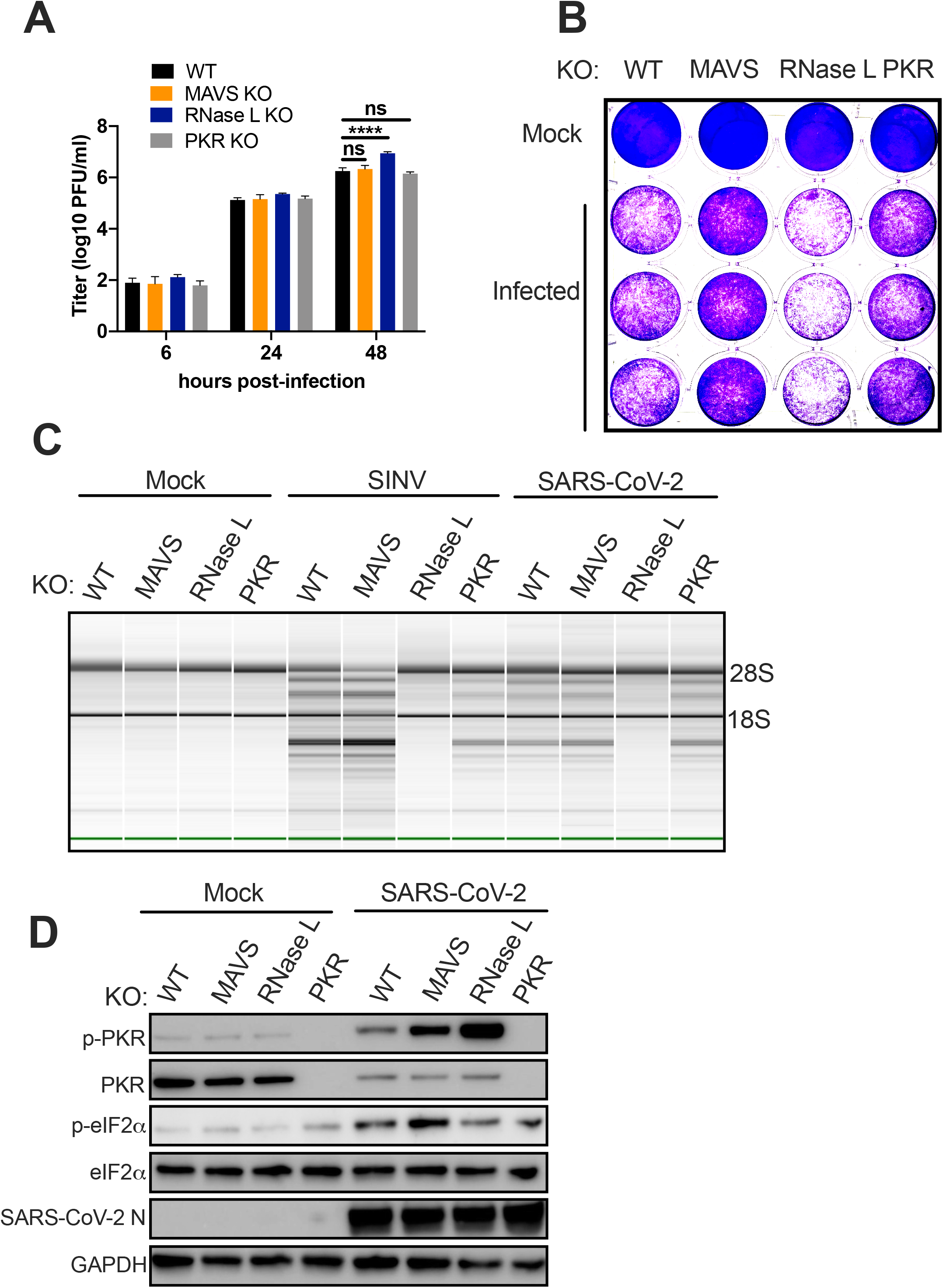
Replication of SARS-CoV-2 is restricted by RNase L independent of PKR or MAVS. Indicated genes were knocked out (KO) from A549^ACE2^ cells using CRISPR-Cas9 engineering. (A) Indicated cell lines were infected with SARS-CoV-2 at MOI=1. At the indicated time points, supernatant was collected and virus quantified by plaque assay on Vero-E6 cells. Values represent mean ± SD (error bars). Statistical significance was determined by two-way ANOVA (****, P < 0.0001; ns = not significant). Data are one representative experiment from at least three independent experiments. (B) Indicated cell lines were mock treated or infected with SARS-CoV-2 at MOI=1. At 48 hpi, cells were fixed with 4% PFA and stained with 1% crystal violet as a marker for live cells. The image is one representative experiment from two independent experiments. (C) The indicated cell lines were mock infected or infected with SARS-CoV-2 or SINV at MOI=1. RNA was harvested 24 hpi (SINV) or 24 and 48 hpi (SARS-CoV-2). Integrity of rRNA was assessed by Bioanalyzer. 28S and 18S rRNA bands are indicated. Data are one representative of two independent experiments. (D) Mock infected or SARS-CoV-2 (MOI=1) infected cells were lysed at 48 hpi and proteins harvested. Proteins were analyzed by immunoblotting using the indicated antibodies. Data are from one representative of two independent experiments. (See also Figure S5).

We assessed rRNA degradation in cells infected with SARS-CoV-2 or SINV (**Fig 9C**) and, as expected, found that rRNA remained intact in the *RNASEL* KO A549^ACE2^ cells, which further validated these cells. However, rRNA was degraded in *PKR* or *MAVS* KO cells, indicating RNase L activation in both of these cell types (**Fig 9C**). Similarly, the PKR pathway was activated by SARS-CoV-2 (**Fig 9D**) and SINV (**Fig S5B**), as evidenced by phosphorylation of PKR and eIF2α, in both *RNASEL* KO and *MAVS* KO cells. More pPKR was detected in *RNASEL* KO cells than WT or *MAVS* KO cells, perhaps due to higher viral titer. Moreover, phosphorylated peIF2α was observed even in absence of PKR, suggesting that at least one other kinase may contribute to phosphorylation of eIF2α during infection with SARS-CoV-2 (**Fig 9D**) but not SINV (**Fig S5B**). These data are consistent with our previous findings that activation of the RNase L pathway does not depend on MAVS signaling in A549 cells infected with SINV or Zika virus (ZIKV) (18, 55), and demonstrate that the PKR pathway can also be activated independently of MAVS. Thus, RNase L and PKR activation occur in parallel with IFN production (**Fig 1**), are not dependent on each other (56).

## Discussion

We evaluated responses to SARS-CoV-2 infection in primary nasal epithelia-derived upper airway cells and iPSC-derived type II airway (iAT2) cells, as well as iPSC-derived cardiomyocytes (iCM), another likely target of infection (32). To complement these studies, we used two lung derived transformed cell lines, Calu-3 cells and two different A549^ACE2^ clones, to more mechanistically dissect activation and antagonism of these pathways by SARS-CoV-2. We found that the extent of IFN induction and signaling is variable among the primary cell types and cell lines used, but is consistently only poorly induced. Interestingly, we show that SARS-CoV-2 infection results in more IFN signaling (phosphorylation of STAT1 and IFN/ISG expression) when compared to MERS-CoV in Calu-3 cells. We also found that SARS-CoV-2 activates RNase L and PKR in both cell lines used, and PKR in iAT2 cells and iCM, but not in primary nasal cells. Using KO cell lines, we demonstrate that RNase L expression significantly impacts SARS-CoV-2 viral titers and CPE observed during infection. These data suggest that while SARS-CoV-2 is generally a weak activator of IFN signaling responses of the respiratory and cardiovascular systems, SARS-CoV-2 can induce the PKR and OAS-RNase L pathways and thus is less adept at antagonizing host responses than MERS-CoV.

As nasal cells are the initial replication site of SARS-CoV-2 and MERS-CoV, we quantified virus replication in infected nasal cell culture. We found that SARS-CoV-2 replicates to higher titer than MERS-CoV, and that the time period for shedding of virus is much longer (**Fig 2A**). We suggest that this longer period of replication in nasal cells and stronger immune responses in Calu-3 cells may in part explain why SARS-CoV-2 is less virulent, yet more contagious than MERS-CoV. Indeed for SARS-CoV-2, R_0_=5.7 (57) while for MERS-CoV, R_0_=0.45 (58).

Infection of all three primary cell types – nasal cells, iAT2 cells, and iCM – resulted in high levels of SARS-CoV-2 replication, while only iCM exhibited obvious CPE (**Figs 2–4**). Syncytia formation was observed in both A549^ACE2^ and Calu-3 cell lines and IFA staining with viral dsRNA-specific antibody (J2) showed SARS-CoV-2 dsRNA localized to perinuclear areas in A549^ACE2^ and Calu-3 cells, which is typical of coronavirus infection (**Fig 5**). The protein expression level of the SARS-CoV-2 host receptor ACE2 (59–61) in primary cells and Calu-3 cells was either low or undetectable, indicating that high levels of receptor are not necessary for productive infection (**Fig 2–4&S3**). This is similar to previous observations in the murine coronavirus (MHV) system where viral receptor CEACAM1a is very weakly expressed in the mouse brain, a major site of infection, and particularly in neurons, the most frequently infected cells (62).

The canonical IFN production and signaling pathways activated by the sensing of dsRNA, an obligate intermediate in viral genome replication and mRNA transcription, provide a crucial early antiviral response (**Fig 1**). However, the role of IFN responses during coronavirus infection is complex and at times contradictory. While IFNs may contribute to pathogenesis later on in infection, coronaviruses, often prevent these responses early on during infection in both animal models and humans (63–66). Indeed, weak IFN responses have been observed during initial stages of SARS-CoV-2 infection, but IFN produced later may contribute to the strong inflammatory responses and resulting immunopathology observed during SARS-CoV-2 infection cytokine storms (67, 68). Providing further evidence for the role of IFN in influencing coronavirus pathogeneis, genetic defects in IFN signaling or the presence of antibodies against type I IFNs are found in a fraction of individuals with severe COVID-19 (28, 29). Genome wide associations of the *OAS1, OAS2, OAS3* genes as well as the *IFNAR2* receptor subunit gene have also been associated with COVID-19 severity (30).

Antagonism of dsRNA-induced antiviral pathways has been well characterized for lineage a (for example, MHV) (11) and lineage c betacoronaviruses (MERS-CoV and related bat viruses) (10), however there is less known about lineage b betacoronaviruses, including SARS-CoV (2002) and SARS-CoV-2. We and others have previously found that both MHV and MERS-CoV betacoronaviruses induce only minimal type I and type III IFNs, and fail to activate RNase L or PKR pathways (11, 47, 50, 69, 70). We found that SARS-CoV-2, like other betacoronaviruses, induced limited amounts of type I and type III IFN mRNAs, although this was somewhat variable among the cell types examined. Using SINV as a control for robust activation of IFN, we detected low levels of type I and type III IFN mRNA in nasal cell, iAT2 cells, and iCM (**Fig 2–4**). However, we observed higher levels of OAS2, an ISG, relative to SINV in iAT2 cells (**Fig 3D**). As we have observed among murine cells, we saw vastly different levels of basal expression of both IFN and ISG mRNAs among the cell types infected (**Fig S2**) (70–72). It is understood that higher basal levels of innate immune response mRNAs typically result in a lower threshold for activation of corresponding responses. Interestingly, we observed significantly higher basal levels, especially IFN-λ, in (uninfected) nasal cells as compared to iAT2 cells and iCM (**Fig S2A**). As major barrier cells, we speculate that this may be important for protection as these cells are more often exposed to infectious agents in the environment. Indeed, it is well documented that IFN-λ serves as an added defense for epithelial cells, which may perhaps explain some of the differences observed in basal gene expression between nasal cells and iCM (73–75). As previously reported in heart tissue, the iCM expressed undetectable levels of both MAVS and RNase L, (23, 76), which is possibly to protect the heart from excessive inflammation.

In A549^ACE2^ cells, SARS-CoV-2 induced low levels of IFN-λ and IFN-β mRNAs and somewhat higher ISG mRNA by 48 hpi, as compared with SINV (**Fig 6A**). We observed greater increases in IFN induction in Calu-3 compared to A549^ACE2^ (**Fig 7A**), which may be at least partially due to higher basal levels of IFNs in the Calu-3 cells (**Fig S2**). Calu-3 cells were employed to directly compare the host response to SARS-CoV-2 infection with that of MERS-CoV and mutant MERS-CoV-ΔNS4ab, which lacks the NS4a and NS4b proteins that inhibit IFN production and signaling (10, 48, 50). In Calu-3 cells, SARS-CoV-2 induced more IFN mRNA than WT MERS-CoV, approaching the level of MERS-CoV-ΔNS4ab (**Fig 7A**). Furthermore, SARS-CoV-2 induced higher levels of ISG mRNAs than MERS-CoV and, in the case of OAS2, higher than MERS-CoV-ΔNS4ab as well. Consistent with this, in Calu-3 cells SARS-CoV-2 and MERS-CoV-ΔNS4ab, but not WT MERS-CoV, promoted STAT1 phosphorylation (**Fig 7C**), which leads to ISG transcription and antiviral responses. Overall, our results displayed a trend of relatively weak IFN responses induced by SARS-CoV-2 in airway epithelial cells with limited ISG induction, when compared with host responses to viruses from other families. This is in argreement with a recent report demonstrating that the SARS-CoV-2 Orf6 encoded protein blocks STAT1 entry into the nucleus, leading to the relatively weak IFN induction (77). Additionally, our data show that enhanced IFN/ISG responses in Calu-3 cells restrict virus production, while lower host responses in A549^ACE2^ cells correlate with higher viral titers (**Fig 5**). Considering how robust ACE2 expression appears dispensable for infection of some cell types (nasal, iAT2, Calu-3), these data also indicate that stronger innate immune responses may be more effective at restricting SARS-CoV-2 replication than low ACE2 expression level.

We found that SARS-CoV-2 was unable to prevent activation of RNase L and PKR, although to different extents among the cell types, unlike MHV and MERS-CoV, which shut down these pathways (10, 11, 69). We observed PKR activation as indicated by phosphorylation of PKR and eIF2α in SARS-CoV-2 infected iAT2 (**Fig 3C**) and iCM (one/two experiments) (**Fig 4C**), but not in nasal cells (**Fig 2C**). However, we did not detect rRNA degradation indicative of RNase L activation in these cell types (**Fig 2E, 3E, 4E**). Activation of both RNase L and PKR were observed in A549^ACE2^ and Calu-3 cells during infection with SARS-CoV-2 (**Fig 8**). In Calu-3 cells, this contrasted MERS-CoV and was more similar to MERS-CoV-ΔNS4ab. Previous studies have shown that MERS-CoV NS4a restricts phosphorylation of PKR by binding dsRNA, reducing its accessibility to PKR (10, 50). Additionally, MERS-CoV NS4b, a 2’-5’ phosphodiesterase, prevents RNase L activation by degrading 2-5A, the small molecular activator of RNase L (10, 47). Current understanding of SARS-CoV-2 protein function infers an absence of these types of protein antagonists, therefore it is not surprising that both of these pathways are activated during infection of both A549^ACE2^ and Calu-3. Indeed, MERS-CoV-ΔNS4ab attenuation compared to WT MERS-CoV, as well as lower SARS-CoV-2 titers than those of MERS-CoV (**Fig 5B**), may be at least in part due to RNase L and PKR activation in addition to IFN/ISG induction in Calu-3 cells.

We found that SARS-CoV-2 did not activate dsRNA-induced pathway responses as robustly as SINV (18, 19), which may be due to CoV antagonists encoded by the nsp genes of the replicase locus (3, 78–80). Most notably, nsp15 encodes an endoribonuclease (EndoU) that has been shown in the MHV system to restrict dsRNA accumulation and thus limit activation of both RNase L and PKR (80, 81). Nevertheless, increased, albeit modest, replication and enhanced cell death in SARS-CoV-2 infected *RNASEL* KO cells indicates that this pathway is activated and indeed restricts replication and downstream cell death caused by SARS-CoV-2 infection (**Fig 9A&B**). In contrast, we found that *PKR* KO had no effect on viral titer and infected cells still produced detectable levels of peIF2α. These results mirror a previous report on SARS-CoV, which found that both PKR and <u>P</u>KR-like <u>ER</u> <u>K</u>inase (PERK) were activated during infection and contributed to eIF2α phosphorylation (82). Our results therefore raise the possibility that SARS-CoV-2 infection activates multiple kinases of the integrated stress response, all of which target eIF2α. We have previously found that MERS-CoV infection inhibits host protein synthesis independent of PKR, so that PKR phosphorylation during MERS-CoV-ΔNS4ab infection did not lead to further reduction (10).

KO of *MAVS* and the consequent loss of IFN production had no significant effect on viral titer or cell death. This is similar to our previous findings demonstrating that RNase L activation can occur independent of virus-induced IFN production during SINV (55) or ZIKV (18) infection in A549 cells, as well as during MHV infection of murine bone marrow-derived macrophages (56). We extend these findings to demonstrate that PKR activation, like OAS-RNase L, can occur independently of MAVS signaling, perhaps explaining the phosphorylation of PKR and eIF2α in iCM, which express undetectable levels of MAVS protein (**Fig 4**). This underscores the importance of the RNase L and PKR antiviral pathways, which can be activated early in infection upon concurrent dsRNA sensing by OAS, PKR, and MDA5 receptors before IFN is produced. Alternatively, these pathways can be activated in cells infected by virus that produce low levels of IFN only late in infection, as we observe here with SARS-CoV-2. Further studies are required to determine whether activation of PKR or RNase L during SARS-CoV-2 infection results in functional outcomes characteristic of these pathways, including inhibition of protein synthesis, induction of apoptosis, cleavage of viral RNA, or induction of inflammatory responses (**Fig 1**). Interestingly, we observed possible RNase L-induced apoptosis in the SARS-CoV-2 infected A549^ACE2^ WT, *MAVS* KO, and *PKR* KO cells, when compared with mock infected counterparts (**Fig 9C**). However, *RNASEL* KO cells displayed the most cell death among the four cell lines, suggesting that virus-induced cell lysis in the *RNASEL* KO cells where viral titers are highest (**Fig 9B**) is more detrimental to cells than RNase L-induced programmed cell death.

We have shown that SARS-CoV-2 activates dsRNA-induced innate immune responses to levels similar to those of a MERS-CoV mutant lacking two accessory proteins that antagonize these pathways, which highlights the distinctions among coronaviruses in interacting with these pathways. However, like MERS-CoV and MHV, SARS-CoV-2 induces limited and late IFN/ISG responses, indicating that proteins antagonizing innate immune responses are likely encoded. Our future studies will focus on identifying specific innate immunity antagonists among lineage b betacoronavirus accessory proteins as well as conserved proteins encoded in the replicase locus.

## Materials and Methods

### Viruses

SARS-CoV-2 (USA-WA1/2020 strain) was obtained from BEI and propagated in Vero-E6 cells. The genome RNA was sequenced was found to be identical to GenBank: MN985325.1. Recombinant MERS-CoV and MERS-CoV-ΔNS4ab were described previously (10) and were propagated in Vero-CCL81 cells. Sindbis virus Girdwood (G100) was obtained from Dr. Mark Heise, University of North Carolina, Chapel Hill (83). Sendai virus (SeV) strain Cantell (84) was obtained from Dr. Carolina B. Lopez (University of Pennsylvania, now Washington University, St Louis). All infections and virus manipulations were conducted in a biosafety level 3 (BSL-3) laboratory using appropriate and approved personal protective equipment and protocols.

### Cell lines

African green monkey kidney Vero cells (E6) or (CCL81) (obtained from ATCC) were cultured in Dulbecco’s modified Eagle’s medium (DMEM; Gibco catalog no. 11965), supplemented with 10% fetal bovine serum (FBS), 100 U/ml of penicillin, 100 μg/ml streptomycin, 50 μg/ml gentamicin, 1mM sodium pyruvate, and 10mM HEPES. Human A549 cells (verified by ATCC) were cultured in RPMI 1640 (Gibco catalog no. 11875) supplemented with 10% FBS, 100 U/ml of penicillin, and 100 μg/ml streptomycin. Human HEK 293T cells were cultured in DMEM supplemented with 10% FBS and 1 mM sodium pyruvate. Human Calu-3 cells (clone HTB-55) were cultured in MEM supplemented with 20% FBS without antibiotics.

### Primary cell cultures

#### Human sinonasal air liquid interface (ALI) cultures

Sinonasal mucosal specimens were acquired from residual clinical material obtained during sinonasal surgery subsequent to approval fromm The University of Pennsylvania Institutional Review Board. ALI cultures were established from enzymatically dissociated human sinonasal epithelial cells (HSEC) as previously described (85, 86) and grown to confluence with bronchial epithelial basal medium (BEBM; Lonza, Alpharetta, GA) supplemented with BEGM Singlequots (Lonza), 100 U/ml penicillin and 0.25 μg /ml amphotericin B for 7 days. Cells were then trypsinized and seeded on porous polyester membranes (2-3× 10^4^ cells per membrane) in cell culture inserts (Transwell-clear, diameter 12 mm, 0.4 μm pores; Corning, Acton, MA). Five days later the culture medium was removed from the upper compartment and the epithelium was allowed to differentiate by using the differentiation medium consisting of 1:1 DMEM (Invitrogen, Grand Island, NY) and BEBM (Lonza), supplemented with BEGM Singlequots (Lonza) with 0.1 nM retinoic acid (Sigma-Aldrich), 100 UI/ml penicillin, 0.25 μg /ml amphotericin B and 2% Nu serum (Corning) in the basal compartment. Cultures were fed every three days for 6 weeks prior to infection with SARS-CoV-2. The day prior infection, the cells were fed and the apical side of the cultures were washed with 100μl of warm PBS X 3.

#### Alveolar organoids and 2D cultures

iPSC (SPC2 iPSC line, clone SPC2-ST-B2, Boston University) derived alveolar epithelial type 2 cells (iAT2) were differentiated and maintained as alveolospheres embedded in 3D Matrigel in CK+DCI media, as previously described (38). iAT2 were passaged_approximately every two weeks_by dissociation into single cells via the sequential application of dispase (2mg/ml, Thermo Fisher Scientific, 17105-04) for 1h at 37°C and 0.05% trypsin (Invitrogen, 25300054) for 15min at 37°C and re-plated at a density of 400 cells/μl of Matrigel (Corning, 356231) in CK+DCI media supplemented with ROCK inhibitor for the first 48h, as previously described (38). For generation of 2D alveolar cells for viral infection, alveolospheres were dispersed into single cells, then plated on pre-coated 1/30 Matrigel plates at a cell density of 125,000 cells/cm2 using CK+DCI media with ROCK inhibitor for the first 48h and then the medium was changed to CK+DCI media at day 3 and infected with SARS-CoV-2 virus.

#### Cardiomyocytes

Experiments involving the use of human iPSCs were approved by the University of Pennsylvania Embryonic Stem Cell Research Oversight Committee. The iPSC line (PENN123i-SV20) used for cardiomyocyte generation was derived by the UPenn iPSC core as previously described (87, 88). This line has been deposited at the WiCell repository (Wicell.org). iPSCs were maintained on Geltrex (Thermofisher Scientific)-coated plates in iPS-Brew XF (Miltenyi Biotec) media at 37°C in 5% CO2/5% O2/90% air humidified atmosphere. Cells were passaged every 5-7 days using Stem-MACS Passaging Solution (Miltenyi Biotec). Differentiation of SV20 into cardiomyocytes (iCMs) was performed using previously described protocols (89, 90). In general, iCMs were >95% positive for cardiac Troponin T staining by FACS. Day 18-25 differentiated cells were replated and used for viral infection experiments.

#### Generation of A549^ACE2^ cells

A549^ACE2^ cells were constructed by lentivirus transduction of *hACE2*. The plasmid encoding the cDNA of *hACE2* was purchased from Addgene. The cDNA was amplified using forward primer 5’-ACTCTAGAATGTCAAGCTCTTCCTGGCTCCTTC-3’ and reverse primer 5’-TTGTCGACTTACGTAGAATCGAGACCGAGGAGAGGGTTAGGGATAGGCTTACCAAAGGAGGTCTGAAC’-3 (contained V5 tag sequences). The fragment containing hACE2-V5 was digested by the XbaI and Sall restriction enzymes from the hACE2 cDNA and was cloned into pLenti-GFP (Addgene) in place of green fluorescent protein (GFP), generating pLenti-hACE2-V5. The resulting plasmids were packaged in lentiviruses pseudotyped with vesicular stomatitis virus glycoprotein G (VSV-G) to establish the gene knock-in cells. Supernatants harvested 48 hours post-transfection were used for transduction into A549 cells. Forty-eight hours after transduction, cells were subjected to hygromycin (1 mg/ml) selection for 3 days and single-cell cloned. Clones were screened for ACE2 expression and susceptibility to SARS-CoV-2 replication.

#### CRISPR/Cas9 engineered cells

*RNASEL, PKR* and *MAVS* KO A549^ACE2^ cells (clone 44) were constructed using the same Lenti-CRISPR system and guide RNA sequences as previously described (19, 54).

#### Viral growth kinetics

The nasal ALI cultures were apically infected with SARS-CoV-2 (MOI=5) or MERS-CoV (MOI=5). Viral stocks were diluted in nasal cell media, 50μl was added to each well, the cells were incubated in 37°C for one hour, then the virus was removed and the cells were wash three times with 200μl of PBS. For viral growth curves, at indicated time points, 200μl of PBS was added to the apical surface, collected 5 minutes later and frozen for subsequent analysis of shed virus by plaque assay. The inserts were transferred to new 24-well plates with fresh media after each collection. For iAT2 or iCM, cells were plated in 12 or 6-well plates, 4X10^5^ cells (iAT2) or 6.25X10^5^ cells per well (iCM), cells were infected with SARS-CoV-2 at MOI=5 (iAT2) or MOI=1 (iCM). At 6, 24, 48 hours postinfection, 200μl of supernatant were harvested and stored in −80°C for infectious virus titration. For infections, cell lines were plated in 12-well plates, A549 and Vero-E6 at 5X10^5^ cells per well and Calu-3 at 3X10^5^ cells per well. Viruses were diluted in serum-free RPMI (A549 infections) or serum-free DMEM (Vero infections) or serum-free MEM (Calu-3 infections) and added to cells for absorption for 1 hour at 37°C. Cells were washed three times with PBS and fed with DMEM or RPMI +2% FBS for Vero and RPMI infections, respectively, or 4% FBS in MEM for Calu-3 infections (47). For virus titration 200μl of supernatant was collected at the times indicated and stored at −80°C for plaque assay on Vero-E6 (SARS-CoV-2) or Vero-CCL81 (MERS-CoV) cells as previously described (91).

#### Plaque assay

Briefly virus supernatant was 10-fold serial diluted and inoculum was absorbed on Vero-E6 cells (SARS-CoV-2) or VeroCCL81 cells (MERS-CoV and MERS-CoV-Δ4ab) for 1 hour at 37°C. Inoculum was overlaid with DMEM plus agarose (either 0.7% or 0.1%) and incubated for 3 days at 37°C. Cells were fixed with 4% paraformaldehyde and stained with 1% crystal violet for counting plaques.

#### Immunofluorescent staining

For nasal ALI culture, following 48 hours of infection, the cultures were fixed in 4% paraformaldehyde at room temperature for 30 minutes. The transwell supports were washed 3 times with PBS prior to excision of the membrane containing the cells. The cells were permeabilized with 0.2% Triton X-100 in PBS and then immersed in PBS with 0.2% Triton X-100, 10% normal donkey serum, and 1% BSA for 60 min at room temperature. Primary antibody incubation was incubated overnight at 4°C (Type IV tubulin, Abcam ab11315, rabbit anti SARS-CoV-2 Nucleocapsid protein, GeneTex, Irvine, CA). Visualization was carried out with Alexa Fluor^®^-conjugated donkey anti-mouse or anti-rabbit IgGs (Thermo-Fisher) (1:1000; 60 min incubation at room temperature). Confocal images were acquired with an Olympus Fluoview System (Z-axis step 0.5μm; sequential scanning). For iAT2, the cell monolayer was fixed using 4% paraformaldehyde (PFA) for 30min, 1X PBS was used to removed PFA and proceed with antibody staining. Fixed cells were treated with a blocking solution containing 0.1% Triton X-100 and 5% donkey serum in 1X PBS for 30min. Immunostaining was performed for SARS-CoV-2 nucleocapsid protein expression using the SARS-CoV-2 nucleocapsid antibody at 1:1000 dilution in blocking solution incubated for 30min. After washing primary antibody away, a secondary Alexa Fluor 488^®^-conjugated donkey anti-rabbit IgG (H+L) antibody(Thermo-Fisher) was used at 1:400 dilution in blocking solution and incubated for 30min. Secondary antibody was washed away with 1X PBS and DAPI was used for nuclear staining at 2.5μg/ml. iCM were fixed in 4% paraformaldehyde and permeabilized with 0.1% Triton X-100 for 15 min. Cells were blocked with 10% normal donkey serum (Sigma D9663) in 0.2% Tween 20 (Biorad 170-6531) for 1hr. Antibodies against cardiac troponin T (cTnT, Abcam ab8295; 1:100 mouse) and SARS-CoV-2 nucleocapsid were incubated with cells in blocking solution overnight at 4 °C. Donkey anti-mouse Alexa Fluor 647^®^-conjugated (Invitrogen A31571) and Donkey anti-rabbit Alexa Fluor 488^®^-conjugated (Invitrogen A21206) were diluted 1:250 in blocking solution and incubated with cells for 2hr at RT. Slides were mounted in Slowfade Gold anti-fade reagent with DAPI (Invitrogen S36939). Images were acquired with BZ-X710 all-in-one fluorescence microscope equipped with BZ-X Viewer software (Keyence Corporation). At the indicated times post-infection cells were fixed onto glass coverslips (Calu-3 coverslips were coated with rat tail collagen type-1: Cell Applications, Inc. Cat. # 122-20) with 4% paraformaldehyde for 30 minutes at room temperature. Cells were then washed three times with PBS and permeabilized for 10 minutes with PBS+0.1% Triton-X100. Cells were then blocked in PBS and 3% BSA for 30-60 minutes at room temperature. Primary antibodies were diluted in blocking buffer and incubated on a rocker at room temperature for one hour. Cells were washed three times with blocking buffer and then incubated rocking at room temperature for 60 minutes with secondary antibodies diluted in blocking buffer. Finally, cells were washed twice with blocking buffer and once with PBS, and nuclei stained with DAPI diluted in PBS (2ng/uL final concentration). SARS-CoV-2 nucleoprotein and dsRNA (J2,1:1000, Scions) were detected. Secondary antibodies were from Invitrogen: goat anti-mouse IgG Alexa Fluor 594^®^-conjugated (A-11005) for J2 and goat anti-rabbit IgG Alexa Fluor 488^®^-conjugated (A-11070) for nucleocapsid. Coverslips were mounted onto slides for analysis by widefield microscopy with Nikon Eclipse Ti2 using a Nikon 40x/0.95NA Plan APO objective and NikonDS-Qi1Mc-U3 12 bit camera. Images were processed using Fiji/Image J software.

#### Western immunoblotting

Cells were washed once with ice-cold PBS and lysates harvested at the indicated times post infection with lysis buffer (1% NP-40, 2mM EDTA, 10% glycerol, 150mM NaCl, 50mM Tris HCl) supplemented with protease inhibitors (Roche – complete mini EDTA-free protease inhibitor) and phosphatase inhibitors (Roche – PhosStop easy pack). After 5 minutes lysates were harvested, incubated on ice for 20 minutes, centrifuged for 20 minutes at 4°C and supernatants mixed 3:1 with 4x Laemmli sample buffer. Samples were heated at 95°C for 5 minutes, then separated on 4-15% SDS-PAGE, and transferred to polyvinylidene difluoride (PVDF) membranes. Blots were blocked with 5% nonfat milk or 5% BSA and probed with antibodies (table below) diluted in the same block buffer. Primary antibodies were incubated overnight at 4°C or for 1 hour at room temperature. All secondary antibody incubation steps were done for 1 hour at room temperature. Blots were visualized using Thermo Scientific SuperSignal west chemiluminescent substrates (Cat #: 34095 or 34080). Blots were probed sequentially with antibodies and in between antibody treatments stripped using Thermo Scientific Restore western blot stripping buffer (Cat #: 21059).

| Primary Antibody | Antibody species | Blocking buffer | Dilution | Catalog number |
| --- | --- | --- | --- | --- |
| pPKR (phospho-T446) [E120] | rabbit | 5% milk/TBST | 1 : 1000 | Abcam 32036 |
| PKR (D7F7) | rabbit | 5% milk/TBST | 1:1000 | Cell Signaling Technology 12297S |
| peif2α (S51) | rabbit | 5% BSA/TBST | 1:1000 | Cell Signaling Technology 9721S |
| eif2α | rabbit | 5% BSA/TBST | 1:1000 | Cell Signaling Technology 9722S |
| GAPDH (14C10) | rabbit | 5% milk/TBST | 1:2000 | Cell Signaling Technology 2118S |
| SARS-CoV-2 N | rabbit | 5% milk/TBST | 1:2000 | GTX135357 (Gentex) |
| MERS-CoV N | mouse | 5% milk/TBST | 1:2000 | 40068-MM10 (Sino Biological) |
| pSTAT1 (Tyr701) | rabbit | 5% BSA/TBST | 1:1000 | Cell Signaling Technology 7649 |
| STAT1 | mouse | 5% BSA/TBST | 1:1000 | Santa Cruz (C136): SC-464 |
| ACE2 | rabbit | 5% milk/TBST | 1:1000 | Cell Signaling Technology 4355S |
| MAVS | rabbit | 5% milk/TBST | 1:1000 | Cell Signaling Technology 24930S |
| V5 | rabbit | 5% milk/TBST | 1:1000 | Cell Signaling Technology 13202S |
| RNase L | mouse | 5% milk/TBST | 1:1000 | Robert Silverman laboratory (Cleveland Clinic) (20) |
| MDA5 | rabbit | 5% milk/TBST | 1:1000 | Cell Signaling Technology 5321S |
| **Secondary Antibody** |  |  |  |  |
| goat anti-rabbit IgG | HRP linked | same as primary | 1:3000 | Cell Signaling Technology 7074S |
| goat anti-mouse IgG | HRP linked | same as primary | 1:3000 | Cell Signaling Technology 7076S |

### Quantitative PCR (RT-qPCR)

A549, Calu-3, and iAT2 cells were lysed at indicated times post infection in RLT buffer and DNase-treated before total RNA was extracted using the RNeasy Plus Mini Kit (Qiagen). RNA from iCM and nasal cells was extracted using TRIzol-LS (Ambion), and DNase-treated using the DNA-*free*^™^ Kit (Invitrogen). RNA was reverse transcribed into cDNA with a High Capacity cDNA Reverse Transcriptase Kit (Applied Biosystems). cDNA was amplified using specific RT-qPCR primers (see Table below), iQ^™^ SYBR^®^ Green Supermix (Bio-Rad), and the QuantStudio^™^ 3 PCR system (Thermo Fisher). Host gene expression displayed as fold change over mock-infected samples was generated by first normalizing cycle threshold (C_T_) values to 18S rRNA to generate ΔC_T_ values (ΔC_T_ = C_T_ gene of interest - C_T_ 18S rRNA). Next, Δ(ΔC_T_) values were determined by subtracting the mock-infected ΔC_T_ values from the virus-infected samples. Technical triplicates were averaged and means displayed using the equation 2^-Δ(ΔCT)^. For basal expression levels, C_T_ values were normalized to 18S rRNA to generate ΔC_T_ values (ΔC_T_ = C_T_ gene of interest - C_T_ 18S rRNA), and displayed as 2^-ΔCt^. Basal expression levels were also calculated as fold change over A549^ACE2^ clone 44 using the equation 2^-Δ(ΔCT)^. Δ(ΔC_T_) values were calculated by subtracting ΔC_T_ values from each cell type from the ΔC_T_ value of A549^ACE2^ clone 44. Absolute quantification of SARS-CoV-2 and MERS-CoV genomes was calculated using a standard curve generated from serially diluted known concentrations of a digested plasmid containing the region of interest. For SARS-CoV-2, construct pcDNA6B-nCoV-NSP12-FLAG encoding the RDRP gene (gift from Dr. George Stark, Cleveland Clinic) was digested with Xho1 and purified by Qiagen QIAquick PCR Purification Kit to be used as a standard in the RT-qPCR reaction. For MERS-CoV, cDNA MERS-D1 (91) containing basepairs 12259–15470 of the MERS-CoV genome was digested with BgII and purified by Qiagen QIAquick PCR Purification Kit to be used as a standard in the RT-PCR reaction. Copy numbers were generated by standard curve analysis in the QuantStudio^™^ 3 software, and copy numbers per ug RNA were calculated based on the volume of cDNA used in the qPCR reaction, and concentration of RNA used to generated cDNA. Primer sequences are as follows:

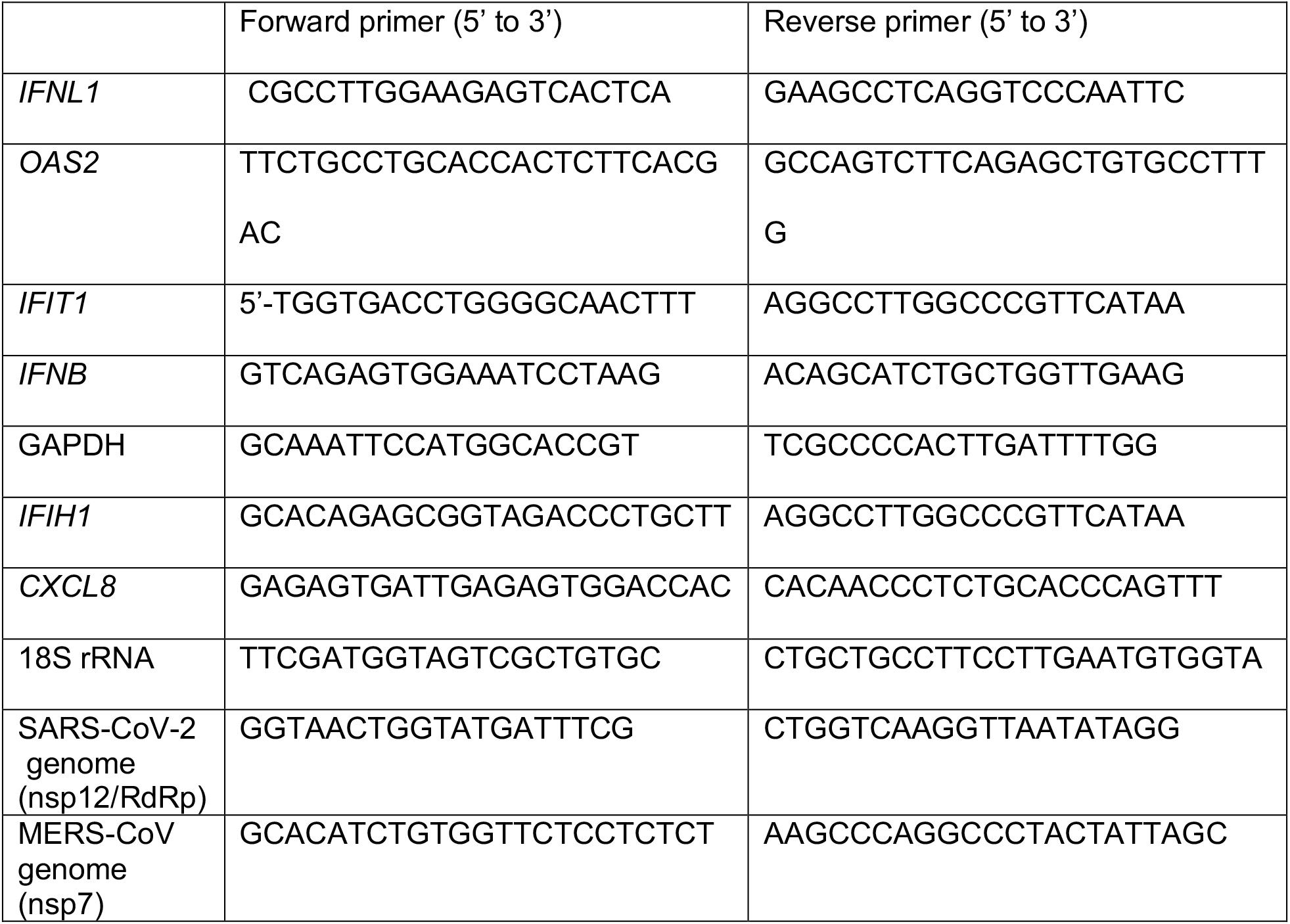

#### Analyses of RNase L-mediated rRNA degradation

RNA was harvested with buffer RLT (Qiagen RNeasy #74106) or Trizol-LS (Ambion) and analyzed on an RNA chip with an Agilent Bioanalyzer using the Agilent RNA 6000 Nano Kit and its prescribed protocol as we have described previously (Cat #: 5067-1511).

#### Statistical analysis

All statistical analyses and plotting of data were performed using GraphPad Prism software (GraphPad Software, Inc., CA). SARS-CoV-2 and MERS-CoV replication trends in nasal cells were analyzed by two-way ANOVA comparing averaged titers from all four donor cells for each virus at each timepoint. MERS-CoV and MERS-CoV-ΔNS4ab viral replication and primary cell RT-qPCR gene expression between SARS-CoV-2 and SINV were analyzed by paired Student *t* test. RT-qPCR analysis in A549^ACE2^ cells was analyzed by one-way ANOVA, comparing SARS-CoV-2 at each timepoint to SINV. RT-qPCR analysis in Calu-3 cells was analyzed by two-way ANOVA, comparing SARS-CoV-2 at each timepoint to MERS-CoV and MERS-CoV-ΔNS4ab. SARS-CoV-2 replication in A549^ACE2^ WT cells compared with A549^ACE2^ KO cells was analyzed by two-way ANOVA. Displayed significance is determined by p-value (P), where * = P < 0.05; ** = P < 0.01; *** = P < 0.001; **** = P < 0.0001; ns = not significant.

## Acknowledgements

We thank Nicholas Parenti for technical help and Dr. Nikki Tanneti for reading the manuscript. This work was supported by NIH grants AI140442 and supplement for SARS-CoV-2 (SRW), AI104887 (SRW and RHS); funds from Penn Center for Coronavirus Research and Other Emerging Pathogens (SRW and YL); NIH grants U01HL148857, R01HL087825, U01HL134745 and R01HL132999 (EM); VA administration grant CX001617 (NAC); NIH grants U01TR001810, N01 75N92020C00005, R01HL095993, and an Evergrande MassCPR award (DNK, JH, and KDA). RT and WY were supported in part by institutional funds from the University of Pennsylania Perelman School of Medicine to the iPSC Core and by NIH grant U01TR001810. DMR was supported in part by T32-AI055400 and CEC was supported in part by T32 NS-007180,

## Author Contributions

Conceptualization: YL, CEC, DMR, SRW

Methodology: YL CEC, DMR, JNW, HMR, WY, NAC, JNP, NDA, MAK, EM, RHS, SRW.

Investigation, Performed experiments: YL, CEC, DMR, JNW, HMR, FLC-D, RT, LHT, BD

Writing–Original Draft: SRW, JNW

Writing–Review & Editing: YL, CEC, DMR, JNW, HMR, SRW, EM, NAC, WY, DNK

Funding Acquisition: SRW, WY, EM, NC, RHS, DNK

Resources: WY, NC, EM, RHS, SRW

Supervision: WY, NC, EM, RHS, SRW

---

**Figure S1.**
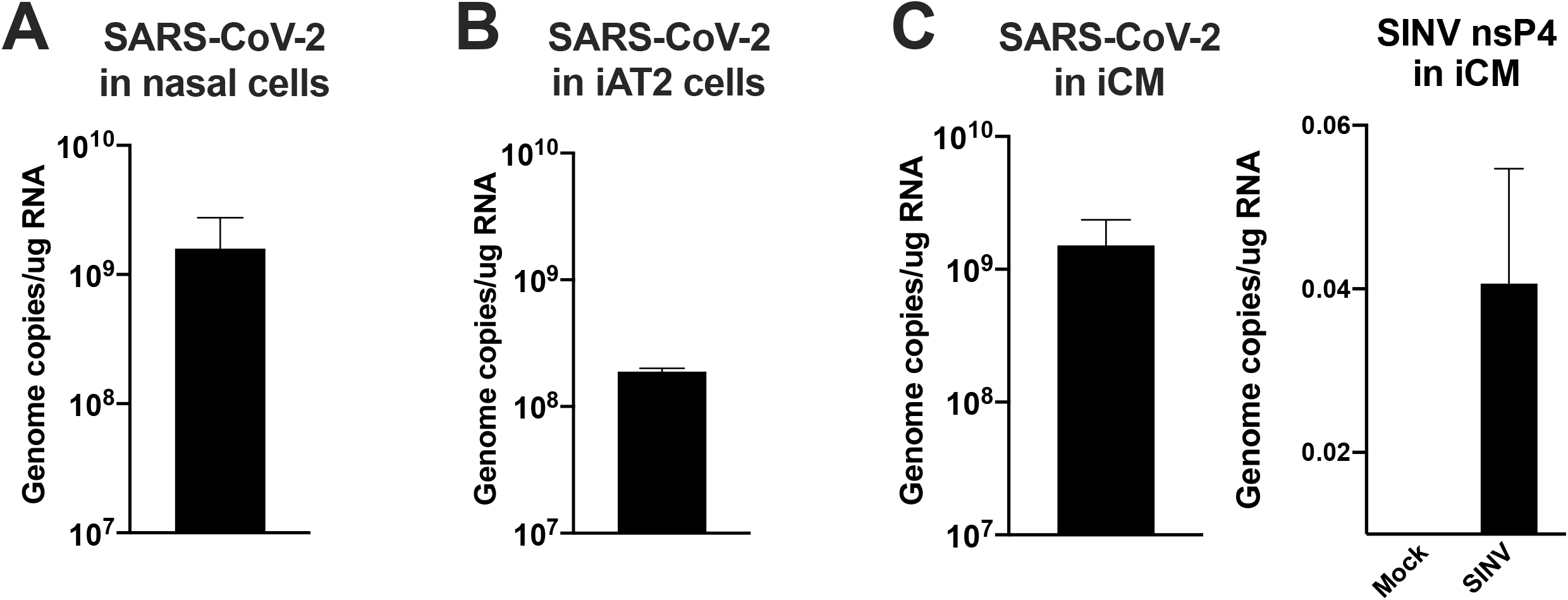
Genome replication in nasal cells, iAT2, and iCM. Nasal (A) and iAT2 cells (B) were infected at MOI=5 with SARS-CoV-2, and (C) iCM at MOI=1 with SARS-CoV-2 or SINV. Total RNA was harvested at 48 hpi (SARS-COV-2) or 16 hpi (SINV) for iAT2 and iCM cells and 120 hpi for nasal cells. Viral genome copies per ug of harvested RNA were calculated by RT-qPCR standard curve generated using a digested plasmid encoding SARS-CoV-2 nsp12. Values are means ± SD (error bars). For SINV (C), cycle threshold (C_T_) values of SINV nsP4 polymerase sequences were normalized to 18S rRNA to generate ΔC_T_ values (ΔC_T_ = C_T_ gene of interest - C_T_ 18S rRNA). Technical triplicates were averaged and displayed using the equation 2^-(ΔCT)^. Data are from one representative experiment of two independent experiments.

**Figure S2.**
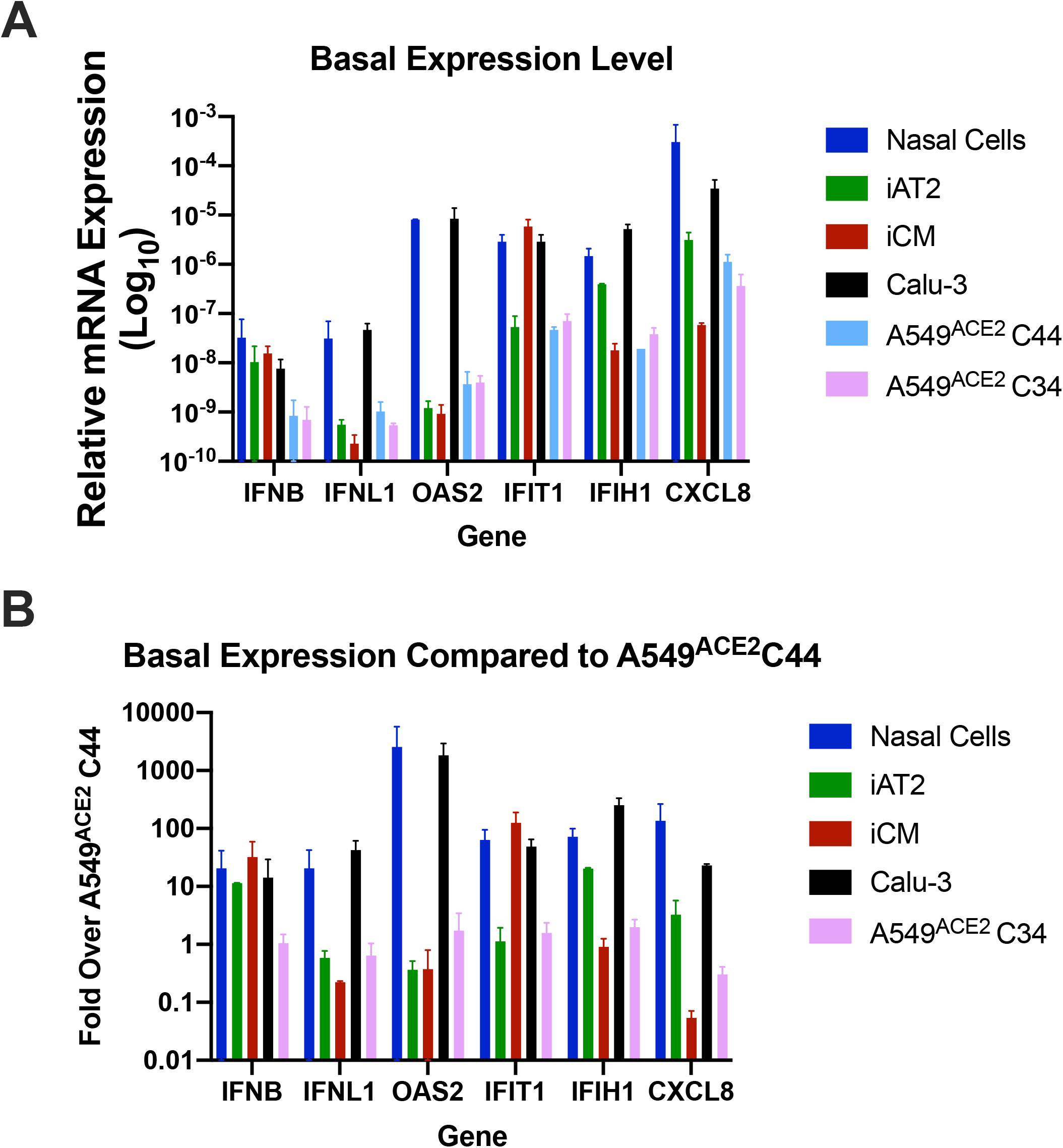
Host basal mRNA expression of uninfected cells. Total RNA was harvested from mock treatment from all indicated cell types after 24 hours incubation. mRNA expression levels of *IFNB, IFNL1, OAS2, IFIT1, IFHI1*, and *CXCL8* were quantified by RT-qPCR. C_T_ values were normalized to 18S rRNA to generate ΔC_T_ values (ΔC_T_ = C_T_ gene of interest - C_T_ 18S rRNA). (A) Basal level of gene expression is displayed for nasal cells, iAT2 and iCM, Calu-3 cells and two clones of A549^ACE2^ cells, displayed as 2^-ΔCt^. (B) Fold expression over A549^ACE2^ C44 values were calculated by subtracting ΔC_T_ values from the indicated cell line from A549^ACE2^ C44 ΔC_T_ values, displayed as 2^-Δ(ΔCT)^. Biological replicates were averaged and values are means ± SD (error bars). Data were generated from at least two independent experiments.

**Figure S3.**
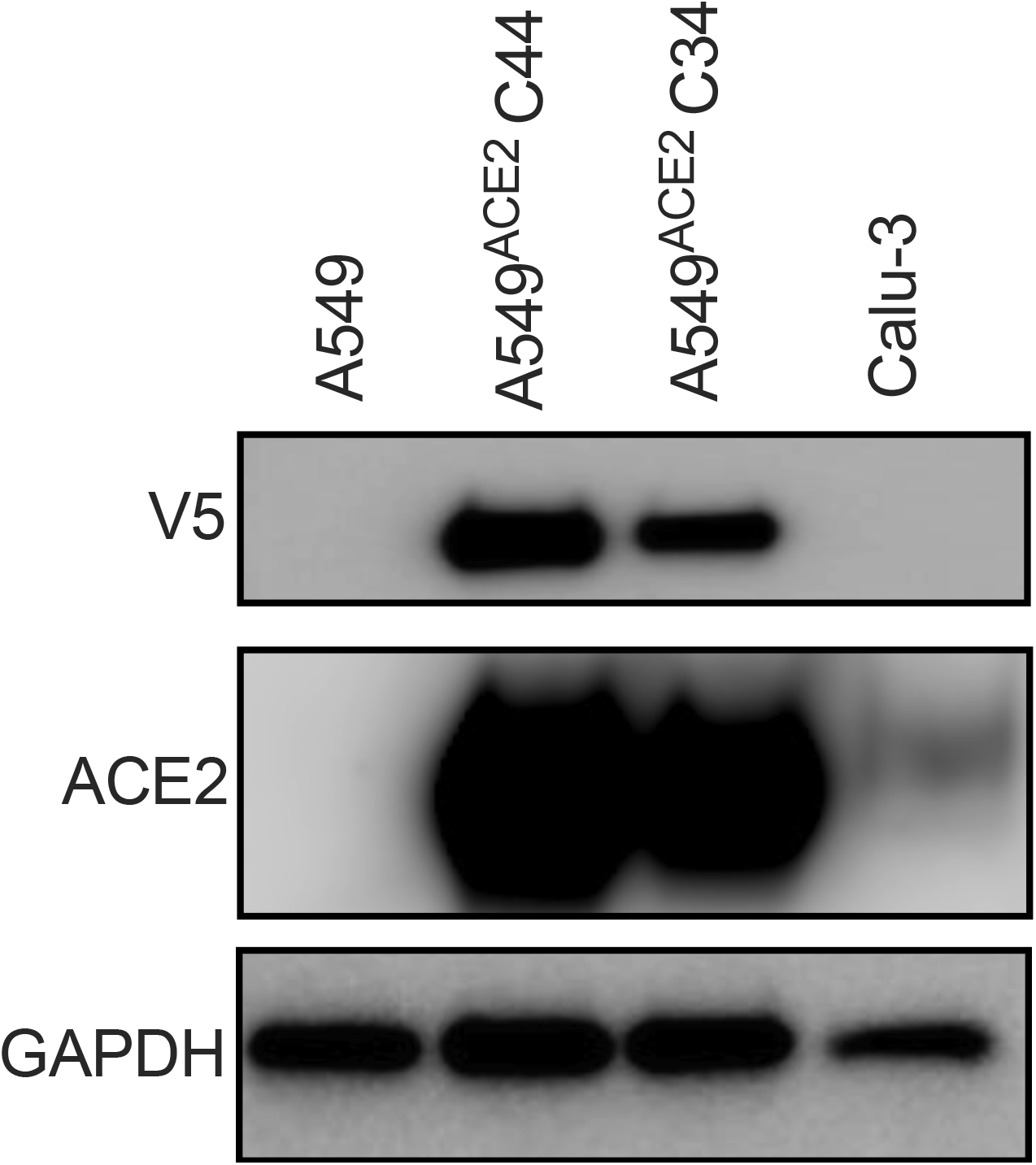
ACE2 protein expression in A549^ACE2^ and Calu-3 cell lines. Parental A549 cells, two A549^ACE2^ clones, and Calu-3 cells were grown in culture before lysis and protein harvest. Protein expression was analyzed by immunoblotting using the indicated antibodies.

**Figure S4.**
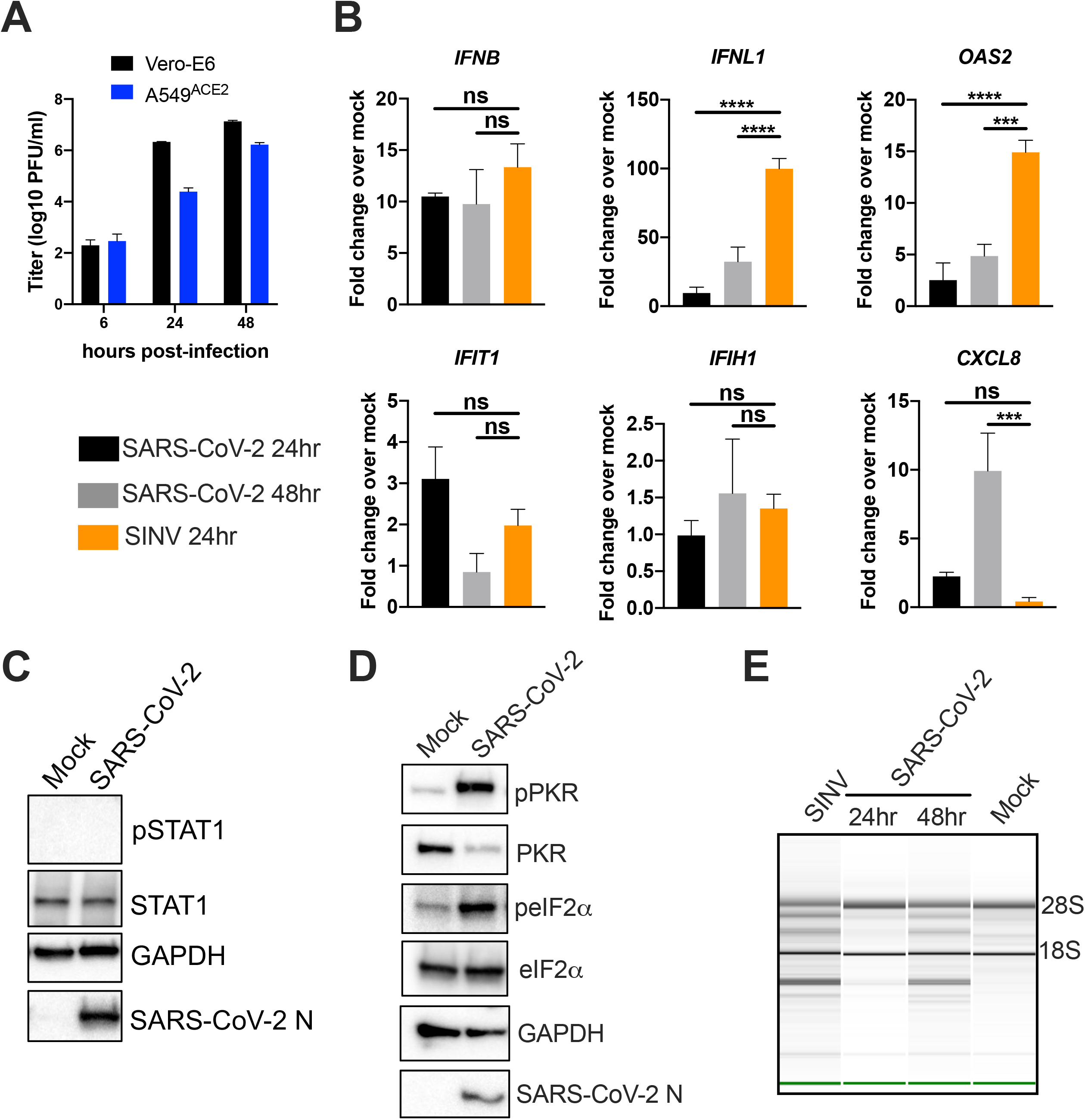
SARS-CoV-2 replication and host responses in a second lung epithelia-derived A549^ACE2^ cell line clone (C34). (A) Vero-E6 or A549^ACE2^ cells were infected with SARS-CoV-2 at MOI=1 and supernatant harvested at indicated times post infection. Infectious virus was quantified by plaque assay on Vero-E6 cells. Values are means ± SD (error bars). (B) A549^ACE2^ cells (C34) were mock infected or infected with SARS-CoV-2 or SINV at MOI=5 and total RNA total RNA harvested at 24 (SINV) or 24 and 48 (SARS-CoV-2) hpi. Expression of *IFNB, IFNL1, OAS2, IFIT1, IFIH1*, and *CXCL8* mRNA was quantified by RT-qPCR. C_T_ values were normalized to 18S rRNA to generate ΔC_T_ values (ΔC_T_ = C_T_ gene of interest - C_T_ 18S rRNA). Fold change over mock values were calculated by subtracting mock infected ΔC_T_ values from virus infected ΔC_T_ values, displayed as 2^-Δ(ΔCt)^. Statistical significance for each gene was determined by one-way ANOVA (***, P < 0.001; ****, P < 0.0001; ns = not significant). Technical replicates were averaged, the means for each replicate displayed, ± SD (error bars). (C&D) A549^ACE2^ cells were infected at MOI=5, lysed at 24 hpi, and proteins harvested for analysis by immunoblotting using the indicated antibodies. (E) A549^ACE2^ cells were infected at MOI=1 (SINV) or MOI=5 (SARS-CoV-2) and total RNA harvested at 24 (SINV) or 24 and 48 (SARS-CoV-2) hpi. Integrity of rRNA was assessed by Bioanalyzer. 28S and 18s rRNA bands are indicated. All data are representative of two or three independent experiments.

**Figure S5.**
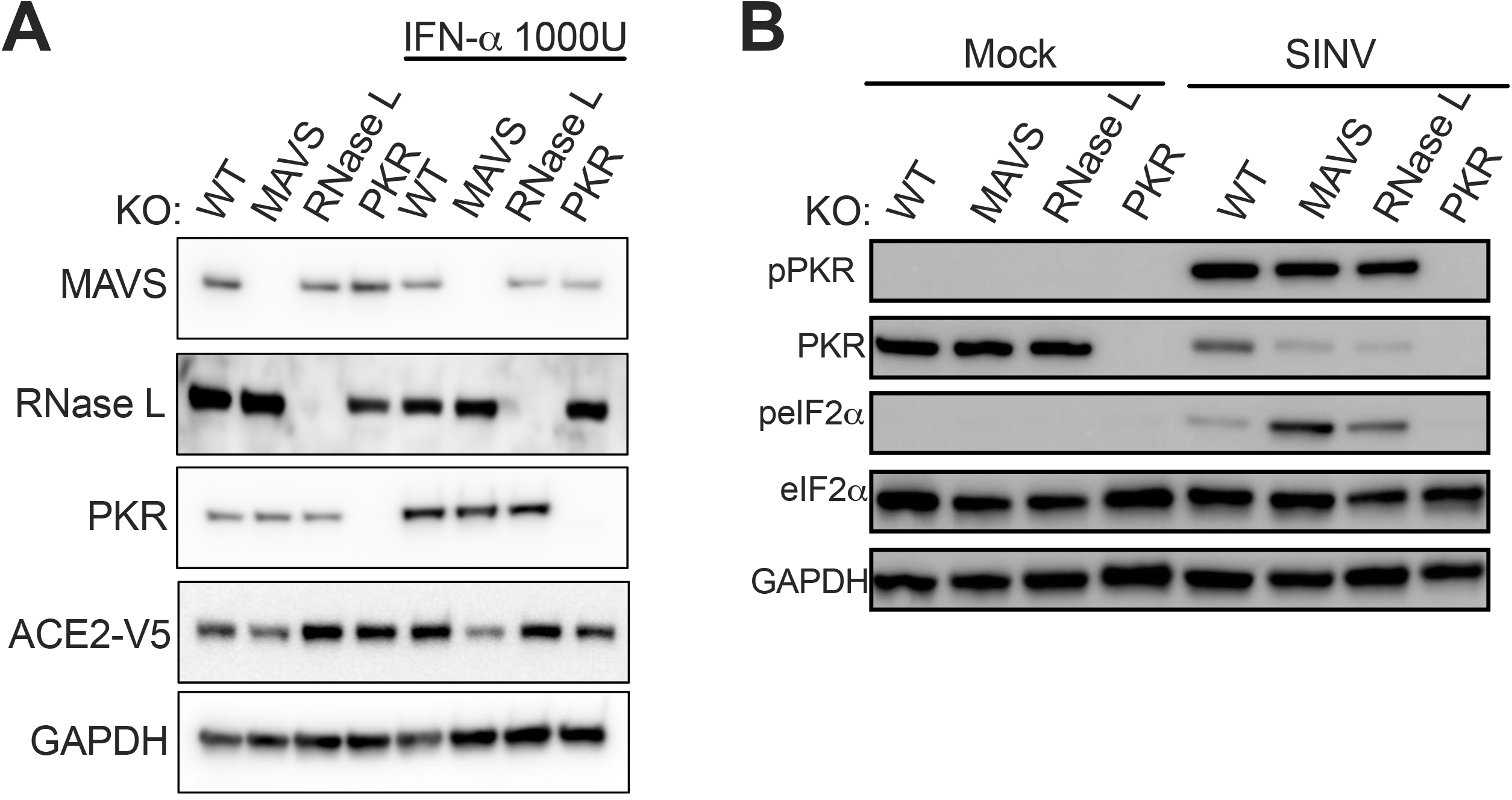
Protein expression in A549^ACE2^ cells. (A) A549^ACE2^ KO cell lines were grown in culture with or without 1000U IFN-α treatment for 24 hours. Cells were lysed and proteins harvested for analysis by immunoblotting using the indicated antibodies. (B) Mock infected or SINV (MOI=1) infected A549^ACE2^ WT or KO cells were lysed at 24 hpi and proteins harvested. Proteins were analyzed by immunoblotting using the indicated antibodies. All data are from one representative of two independent experiments

